# RORγt^+^ B cells express proinflammatory cytokines and promote allo- and auto- immunity

**DOI:** 10.1101/2023.09.22.558524

**Authors:** Qing Ding, Yufan Wu, Adam Zuchowski, Elena Torlai Triglia, Jennifer L. Gommerman, Ayshwarya Subramanian, Vijay K. Kuchroo, David M. Rothstein

## Abstract

B cells can express pro-inflammatory cytokines that promote a wide variety of immune responses. Here we show that B cells expressing the phosphatidylserine receptor TIM-4, preferentially express IL-17A, as well as IL-22, IL-6, IL-1β, and GM-CSF - a pattern of cytokines reminiscent of pathogenic Th17 cells. Expression of this proinflammatory module requires IL-23R signaling and selective expression of RORγt by TIM-4^+^ B cells. TIM-4^+^ B cell-derived-IL-17A not only enhances the severity of experimental autoimmune encephalomyelitis (EAE) and promotes allograft rejection but also acts in an autocrine manner to prevent their conversion into IL-10-expressing B cells with regulatory function. Thus, IL-17A acts as an inflammatory mediator and enforces the proinflammatory activity of TIM-4^+^ B cells. Moreover, TIM-4 serves as a broad marker for RORγt-expressing effector B cells (Beff). These findings allow further study of the signals regulating Beff differentiation and effector molecule expression.

---

In addition to a primary role in humoral immunity, B cells are potent regulators of the immune response through antigen presentation, co-receptor engagement and production of cytokines(*1–6*). Regulatory B cells (Bregs) inhibit immune responses through the expression of suppressive cytokines and coinhibitory molecules(*4–7*). The phosphatidylserine receptor, TIM-1, is both a broad and functional marker for Bregs that play an essential role in restraining tissue inflammation and maintaining self-tolerance(*7, 8*). In addition to being a marker of Breg identity, TIM-1 regulates the expression of a “regulatory module” that includes IL-10 and various coinhibitory molecules including TIGIT(*7*). As such, specific deletion of *Havcr1* (TIM-1 gene) in B cells results in spontaneous systemic autoimmunity characterized by inflammatory infiltration of multiple organs accompanied by dermatitis, weight loss/rectal prolapse, and paralytic neuroinflammatory disease(*7*).

In contrast to Bregs, effector B cells (Beffs), express proinflammatory cytokines that promote anti-microbial responses, autoimmunity, and allograft and tumor rejection(*2, 5, 6, 9*). For example, B cell-derived IL-2 and TNFα are required for clearance of *H. polygyrus* infection(*10*). B cell-derived IL-6 enhances Th1 and Th17 responses and increases the severity of EAE (a murine model of multiple sclerosis), while B cell-derived IFNγ promotes Th1 responses essential for generation of proteoglycan-induced arthritis and promotes extrafollicular versus germinal center responses during acute infection(*11–13*). During *T. cruzi* infection, a pathogen-associated trans-sialidase was reported to specifically induce IL-17A expression in B cells through a RORγt-independent mechanism, promoting pathogen clearance(*14*). However, a broader characterization and role for IL-17A in Beff function has not been elucidated. In humans, proinflammatory cytokines such as GM-CSF, Lymphotoxinα, and TNFα produced by B cells have been implicated in multiple sclerosis and B cell depletion with anti-CD20 reduces T cell hyperreactivity and is now the treatment of choice for primary progressive and relapsing forms of the disease(*15, 16*).

Unlike Bregs, the phenotype of Beffs has not been well-studied, impeding our understanding of their generation and effector functions. We previously showed that TIM-4^+^ B cells are enriched for IFNγ and enhance tumor and allograft rejection – activity opposite to that of TIM-1^+^ Bregs(*9*). However, no more broadly unifying marker for Beffs has been identified and it remains unclear whether B cells expressing different inflammatory cytokines are related in phenotype or are regulated by common signaling mechanisms. It is also unknown whether Bregs and Beffs bear any functional relationship. Here, we characterize the role of TIM-4^+^ B cells as Beffs using high-throughput sequencing and murine models of autoimmunity and transplantation. We show that in addition to IFNγ, TIM-4^+^ B cells express a unique proinflammatory module driven by RORγt expression that includes IL-17A and additional cytokines resembling those expressed by pathogenic Th17 cells. In addition to its proinflammatory role, B cell-derived IL-17A also acts as an autocrine factor that enforces expression of the proinflammatory module by TIM-4^+^ B cells and prevents their conversion into IL-10-expressing B cells with regulatory function. Thus, TIM-4 is a marker that links together B cells expressing most of the proinflammatory cytokines associated with Beff function, and we identify IL-17A as both a B cell proinflammatory cytokine and regulator of Beff versus Breg function.

## RESULTS

### TIM-4^+^ B cells express IL-17A

As previously shown, TIM-4 is expressed on a subset of splenic B cells similar in size but largely distinct from TIM-1^+^ B cells (**Fig. 1A)**(*9*). In addition to IFNγ production, we now show that TIM-4^+^ B cells from alloimmunized mice are also enriched for IL-17A production compared to total (CD19^+^) B cells or TIM-1^+^ B cells (**Fig. 1B-C**), and given their relatively low frequency of IL-10 expression, TIM-4^+^ B cells have a markedly higher IL-17A:IL-10 ratio than TIM-1^+^ B cells (**Fig. 1B-D**). In contrast, a majority of B cells lack both TIM-1 and TIM-4 (double negative; DN) and produce minimal IL-10, IL-17A or IFNγ (**Fig. 1B** and (*9*)). Most B cell-derived IL-17A is produced by the TIM-4^+^ subset, as confirmed by analysis of B cells from alloimmunized IL-17A GFP-reporter mice either in the absence or presence of *in vitro* stimulation with PMA and ionomycin plus brefeldin A (“PIB”) for 5 hours to enhance cytokine expression (**Fig. 1E; Supplementary Fig. 1A**; B cell gating strategy). The frequency of B cells expressing TIM-4 and the frequency of TIM-4+ B cells producing IL-17 was higher in spleen than in LN or bone marrow (**Supplementary Fig. 1D**). Similar levels of IL-17A were observed after immunization of mice for EAE (MOG_35-55_, CFA, and pertussis toxin), suggesting that IL-17A production is a generalized response by a portion of TIM-4^+^ B cells to immunizing stimuli (**Supplementary Fig. 1C, D**).

**Fig 1.**
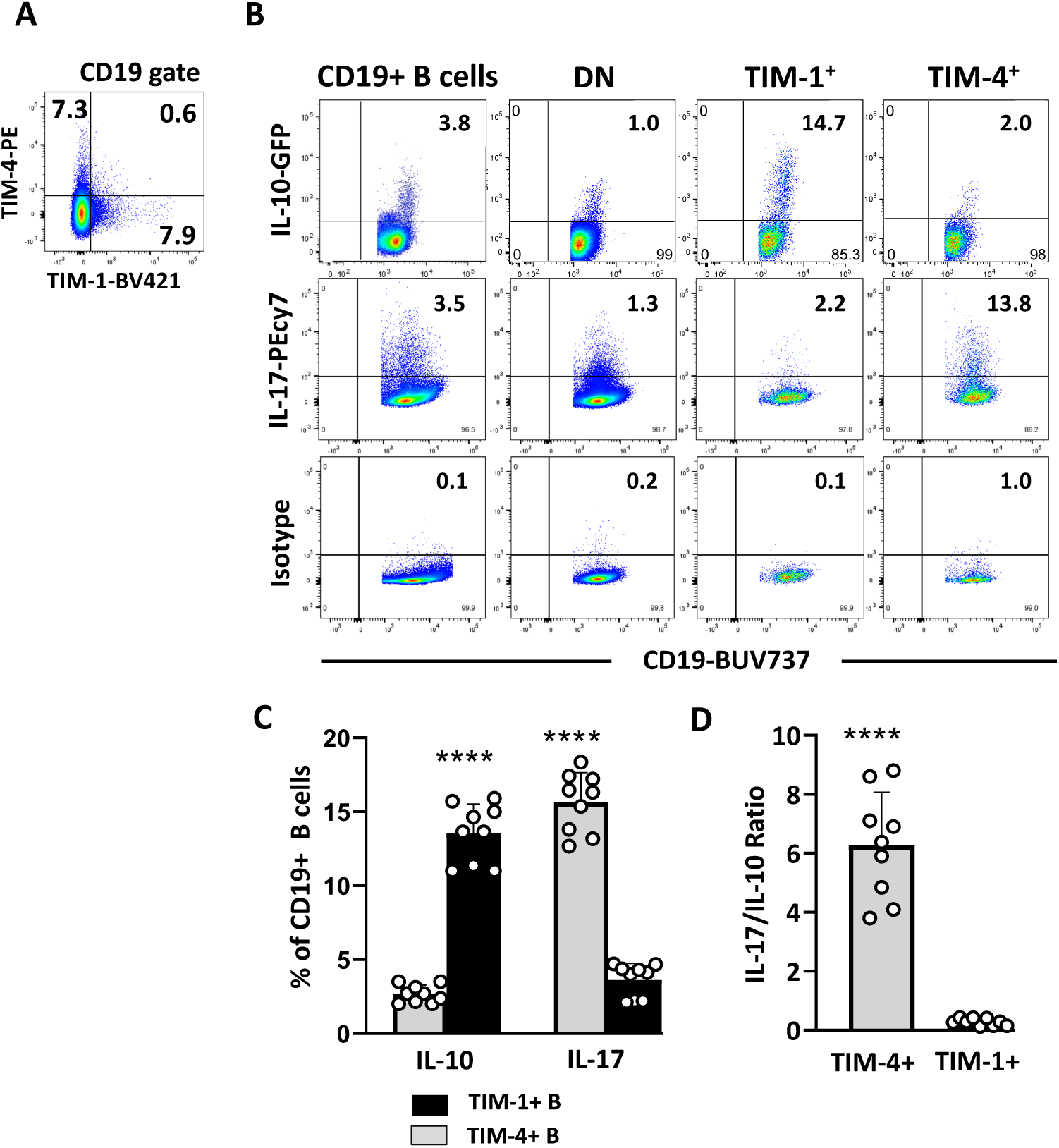
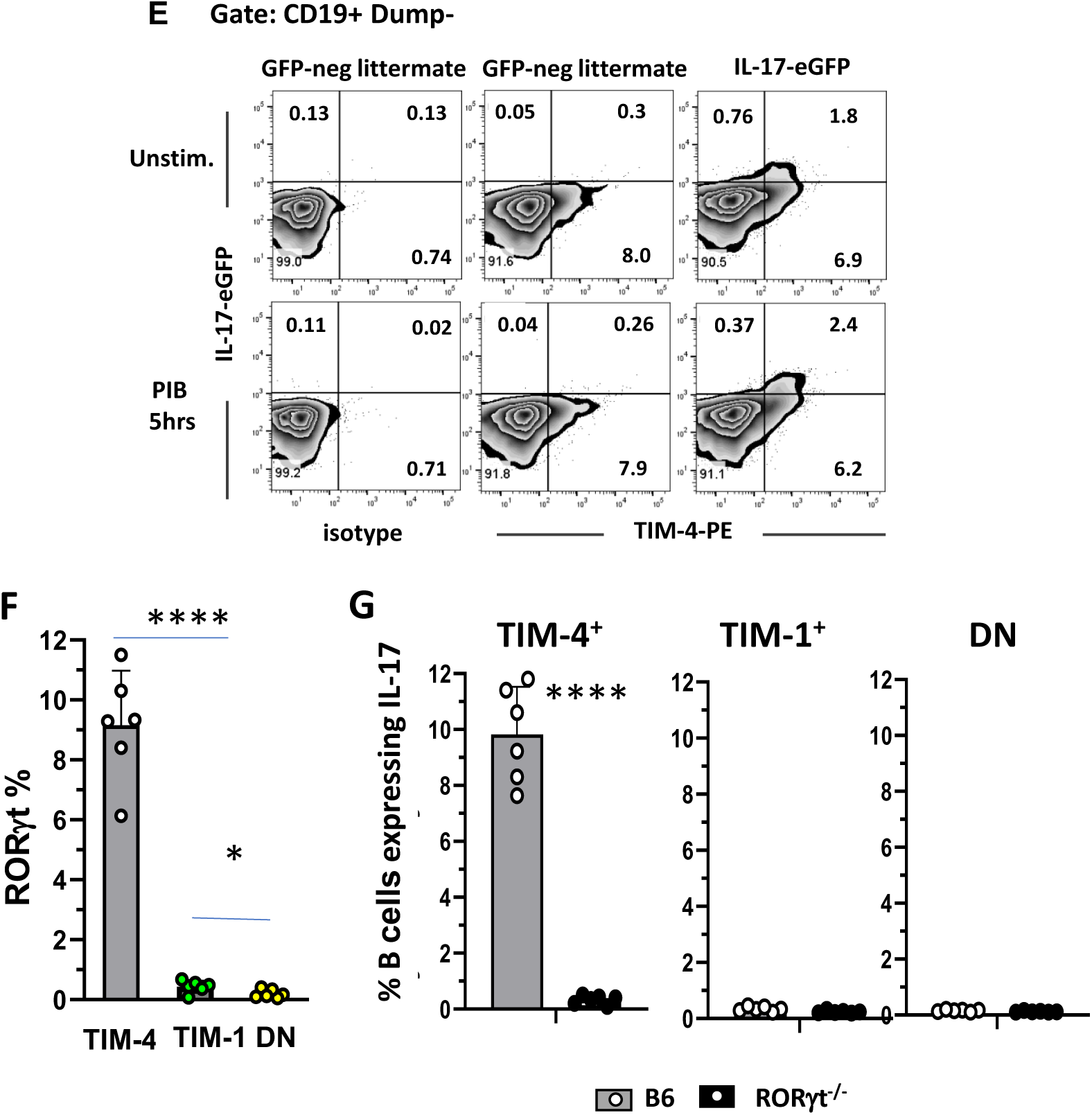
TIM-4^+^ B cells are distinct from those expressing TIM-1^+^ and preferentially express IL-17A. Splenic B cells from C57BL/6 mice 14 days after alloimmunization. (**A, B**) Representative flow cytometry plots: **(A)** TIM-1 and TIM-4 expression on CD19^+^ B cells and **B**) IL-10 and IL-17A expression on total (CD19^+^) B cells, TIM-1^-^TIM-4^-^ (DN), TIM-1^+^, and TIM-4^+^ B cell subpopulations. IL-17A isotype (control) shown in bottom row. (GFP-negative control for IL-10-GFP reporter not shown). (**C**) Bar graph showing mean frequency (+SD) of IL-10 and IL-17A expression on TIM-1^+^ vs. TIM-4^+^ B cells based on flow cytometry, as in (**B**) (n=9 mice per group in 3 individual experiments). (**D**) Bar graph showing the ratio of IL-17A^+^/IL-10^+^ cells within TIM-1^+^ and TIM-4^+^ B cell subsets (mean + SD) based on data in (**C**). (**E**) Representative FACS plots showing TIM-4 and IL-17A-EGFP expression on unstimulated vs. PIB stimulated CD19^+^ B cells from alloimmunized IL-17A-EGFP reporter mice. (n=3 mice per group). (**F**) Bar graph showing the mean frequency (± SD) of RORγt expression in DN, TIM-1⁺, and TIM-4⁺ B-cell subpopulations. (n=6 mice per group in 2 individual experiments) (**G)**. Bar graphs showing the mean frequency (± SD) of IL-17 expression in DN, TIM-1⁺, and TIM-4⁺ B-cell subpopulations from alloimmunized RORγt-knockout and WT B6 mice. n=6 mice per group in 2 individual experiments. Statistics: *p< 0.05 vs. other groups; **** p<0.0001 vs. other groups.

To address why IL-17 expression by B cells is largely limited to the TIM-4+ subset, we examined RORγt expression. We found that RORγt is almost exclusively expressed by TIM-4+ B cells (**Fig. 1F, Supplementary Fig. 1E**). Further, deletion of *Rorc* (encoding RORγt) leads to loss of IL-17 expression by B cells (**Fig. 1G**). Thus, as in T cells and Type-3 ILCs, B cell IL-17 is RORγt dependent. RORγt is expressed by a small portion (<3%) of TIM-4+ B cells in naïve mice but increases to ∼5% after immunization and 9% after 5h *in vitro* stimulation with PIB (**Fig 1F**, **Supplementary Fig. 1F**).

### B cell IL-17A is an important driver of allo- and auto- immune responses

We first examined the role of IL-17A produced by B cells in MOG_35-55_-induced EAE, where Th17 cells and Bregs both play a prominent role(*7, 17*). In agreement with previous studies, μMT mice exhibit severe and unremitting EAE, and this is reduced by the transfer of WT B cells which contain Bregs (**Fig. 2A**)(*17*). Since WT B cells also include IL-17A-producing TIM-4+ Beffs, we hypothesized that transfer of *Il17a*^-/-^ B cells might further reduce EAE severity. Indeed, transfer of *Il17a*^-/-^ B cells into μMT mice reduced EAE severity well below that seen after transfer of WT B cells.

**Fig 2.**
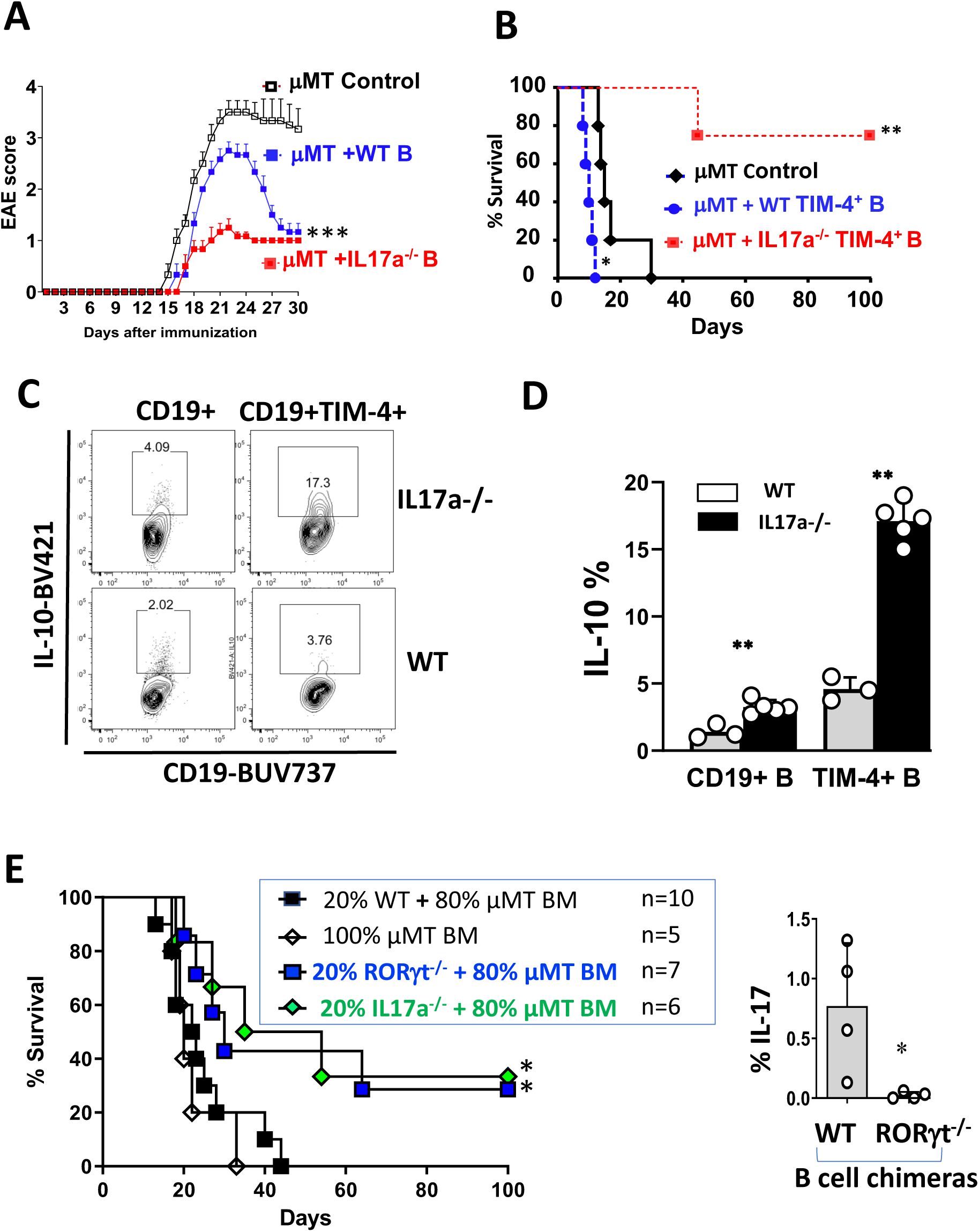
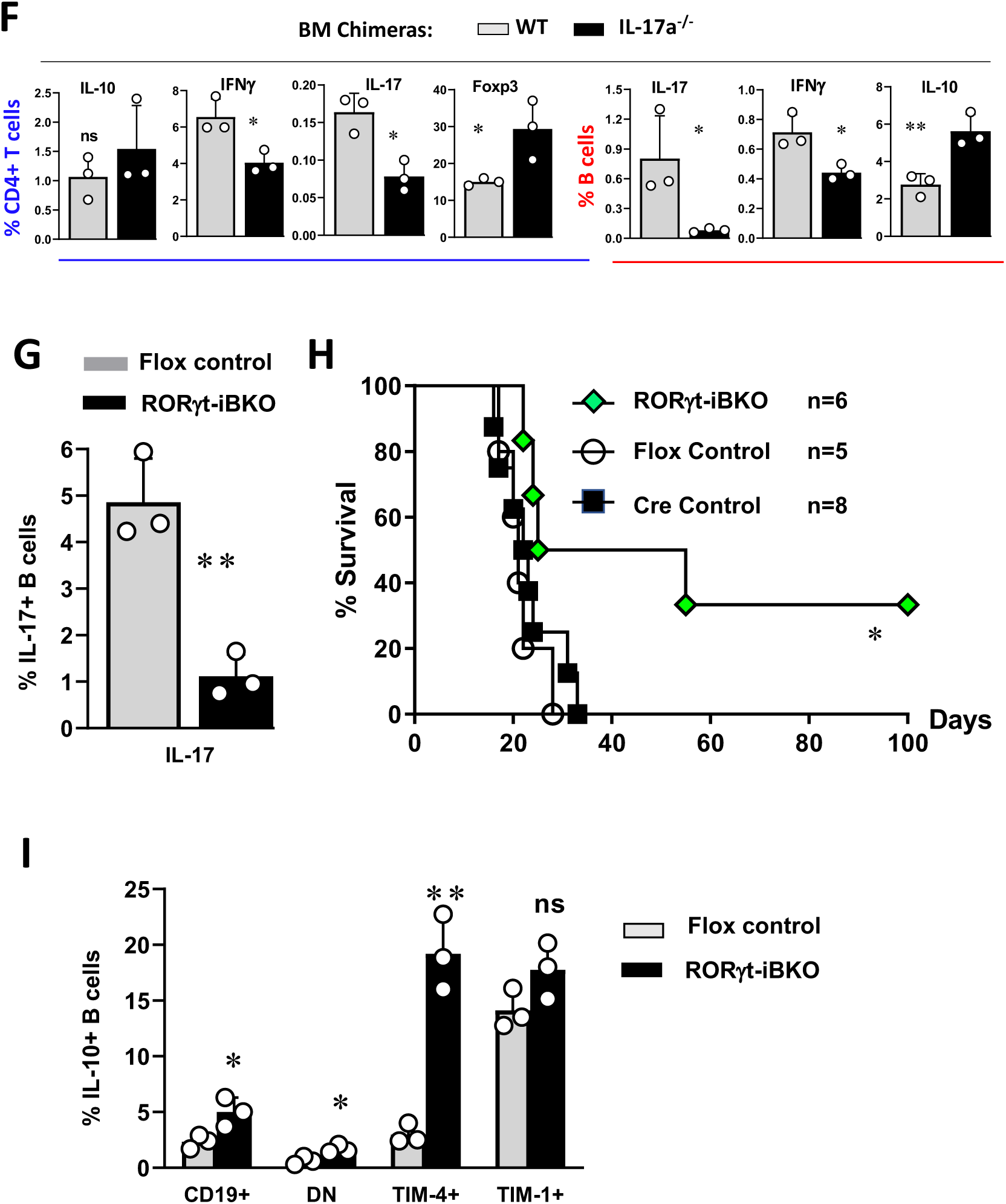
IL-17-deficient B cells are regulatory. **(A)** EAE was induced in µMT mice that received no B cells, or adoptive transfer of 10 X10^6^ B cells from naive WT or IL-17A^-/-^ mice. Fig. shows mean (+SEM) EAE scores over time (n≥6 mice per group; representative of 2 independent experiments). ***p < 0.001 by 2-way ANOVA. (**B**) Kaplan–Meier plots showing survival of BALB/c islet allografts in diabetic µMT (B6) recipients that were untreated (control) or received 5×10^6^ syngeneic TIM-4^+^CD19^+^ B cells from (day 14) alloimmunized WT or IL-17^-/-^ donor mice. n=4-5 mice/group. *p <0.05; **p <0.01 versus control. (**C**) Representative flow cytometry plots of IL-10 expression by CD19^+^ and TIM-4^+^ B cells from alloimmunized IL-17A^-/-^ versus WT B6 mice. **(D**) Bar graph showing mean frequency (+SD) of IL-10 expression on total (CD19^+^) and TIM-4^+^ B cells represented in panel (C). n = 3-5 mice/group. **p < 0.01. **(E)** μMT BM chimeric mice were reconstituted with 100% μMT BM or with 80% μMT BM plus 20% *Il17^−/−^*, *Rorc^−/−^,* or WT BM. **Left panel**: Kaplan-Meier plots showing BALB/c islet allograft survival in diabetic BM chimeric recipients (n=5-10 mice/group) *p < 0.05 vs. Controls. **Right panel**: Quantitation of IL-17A expression (intracellular staining) in alloimmunized WT vs. *Rorc^−/−^* BM chimeras (n=4 mice/group) *p < 0.05. **(F)** WT and *Il17a* BM chimeric mice were alloimmunized and cytokine (IL-10, IL-17, and IFN-γ) and Foxp3 expression on endogenous CD4^+^ T cells and CD19^+^ B cells was determined by intracellular flow cytometry. *p<0.05; **p<0.01. **(G)** Quantitation of frequency of IL-17^+^ B cells in RORγt-iBKO vs. Rorc^fl/fl^ (Flox Control) mice 14 days after alloimmunization. ** p<0.01. **(H)** Kaplan-Meier plots of BALB/c islet allograft survival in Tamoxifen-treated diabetic RORγt-iBKO versus Flox Control and hCD20.ERT2.Cre^+/-^ (Cre control) recipients. n=5-8 mice/group *p<0.05. **(I)** Bar Graph showing mean frequency (mean + SD) of IL-10 expression on total B cells and B cell subpopulations (n=3 mice per group). *p<0.05, **p < 0.01 RORγt-iBKO vs. Flox Control mice.

We next examined the role of B cell IL-17 production in alloimmunity, which results in a robust innate and polyclonal T cell response, but where the role of type-17 immunity has not been established(*18, 19*). To remove the potentially confounding influence of Bregs from the findings, we transferred sort-purified TIM-4^+^ Beffs from alloimmunized syngeneic (B6) mice into chemically diabetic μMT (B6) mice that received fully H-2 mismatched BALB/c islet allografts but were otherwise untreated. As we previously demonstrated, transfer of WT TIM-4^+^ B cells accelerated islet allograft rejection compared to control μMT recipients without B cell transfer (**Fig. 2B**). Transfer of *Il17a*^-/-^ TIM-4^+^ B cells from alloimmunized mice no longer accelerated allograft rejection, and in fact, markedly prolonged graft survival. In attempts to explain the suppressive activity of TIM-4^+^ B cells from *Il17a*^-/-^ mice, we examined their cytokine expression. Compared to WT, *Il17a*^-/-^ B cells produced IL-10 with twice the normal frequency (**Fig. 2C, D**). In particular, IL-10 frequency within *Il17a*^-/-^ TIM-4^+^ B cells increased 3-fold, now similar to that of WT TIM-1^+^ B cells (**Fig. 1C**).

To ensure that the increased B cell IL-10 observed above was not due to B cell dysregulation arising in globally IL-17A-deficient mice, we generated mixed bone marrow (BM) chimeras whose B cells specifically lacked IL-17 or RORγt expression. These were compared to Control BM chimeras expressing WT B cells or lacking B cells. Chemically diabetic BM chimeras underwent islet transplant surgery as described above. Compared to Control chimeras, those whose B cells specifically lacked either IL-17 or RORγt exhibited significantly prolonged islet allograft survival and ∼30% of these recipients never rejected their allografts (**Fig. 2E**).

We found that endogenous CD4^+^ T cells from the B cell *Il17a*^-/-^ BM chimeras expressed significantly less IL-17A and IFNγ and exhibited a 2-fold increase in Foxp3^+^ Tregs compared to CD4^+^ T cells from BM chimeras with WT B cells (**Fig. 2F**). To examine the impact of B cell-intrinsic IL-17 deficiency on antigen-specific T cells within the context of an alloimmune response, B cell *Il17a*^-/-^ B cell BM chimeras vs. WT B cell control BM chimeras were immunized with Act-mOva.BALB/c.B6 F1 splenocytes, followed by adoptive transfer of CFSE-labelled OTII CD4^+^ T cells(*20, 21*). IL-17A-deficiency in B cells was associated with decreased proliferation (day 4) and altered frequency of cytokine production (day 7) by OTII CD4^+^ T cells. This included a 2-fold decrease in IFNγ, and ≥1.7-fold increase in IL-10 and IL-4 (**Supplementary Fig. 2A**). Similar changes in cytokine production and proliferation were observed in transferred CD8^+^ OT-1 cells (**Supplementary Fig. 2B**). Thus, loss of Beff-derived IL-17A, reduced antigen-specific T cell proliferation and the T cell response was skewed towards less inflammatory and more regulatory activity. As expected, endogenous B cells from alloimmunized *Il17a^-/-^* B cell BM chimeras produced negligible IL-17A and also exhibited a 30% decrease in frequency of IFNγ expression, compared to B cells from WT BM chimeras (**Fig. 2F).** Similar to B cells from global *Il17a-/-* mice, B cells from BM chimeras whose B cells lacked *Il17a expression,* exhibited a two-fold increase in IL-10 frequency. This suggested that intrinsic expression of IL-17 by B cells suppresses their ability to produce IL-10.

In a further attempt to separate B cell IL-17-deficiency from increased IL-10, we generated mice with an inducible knockout of *Rorc* in B cells (*hCd20-ERT2.Cre* X *Rorc*^fl/fl^; RORγt-iBKO). These mice are entirely normal until being placed on Tamoxifen (TAM) chow to acutely induce B cell-specific deletion of RORγt. B cells from TAM-treated RORγt-iBKO mice exhibited an 80% reduction in B cell-derived IL-17A compared to those from *Rorc*^fl/fl^ (Flox control) mice (**Fig. 2G)**. RORγt-iBKO mice exhibited prolonged islet allograft survival compared to either Flox control or *hCd20-ERT2.Cre* (Cre control) mice, with 33% surviving long-term (**Fig. 2H**), very similar to that seen in the B cell *Il17a*^-/-^ or *Rorc*-/- BM chimeras above. Furthermore, iRORγt-iBKO mice exhibited slower onset and decreased peak and chronic phase EAE severity (**Supplementary Fig. 2C**). Analysis of B cells from alloimmunized RORγt-iBKO mice again demonstrated increased frequency of IL-10 production, ∼2-fold by total B cells and 5-fold by TIM-4^+^ B cells (**Fig. 2I and Supplementary Fig. 2D**). Freshly isolated TIM-1^+^ B cells (which normally express little RORγt), showed no significant increase in IL-10 with *Rorc* deletion. Thus, even the acute loss of RORγt and IL-17A in developmentally normal mice results in dysregulated IL-10 production in the TIM-4^+^ Beff subset. Furthermore, the loss of proinflammatory IL-17A and increased IL-10 production by B cells results in increased allograft survival and decreased EAE severity.

### Regulation of IL-17A and IL-10 expression in B cells by IL-23, IL-17, and RORγt

Given the aberrant production of IL-10 by TIM-4+ B cells lacking RORγt or IL-17, we further studied the regulation of IL-17 and IL-10 expression by sort-purified B cells *in vitro*. Stimulation with anti-IgM for 24 hours maintained the frequency of IL-17A production similar to that observed in freshly isolated CD19+ B cells (**Supplementary Fig. 3A** compared to **Figs. 1B, E**). Culture with IL-23 alone increased IL-17 frequency by 1.7-fold, whereas stimulation with anti-IgM plus IL-23 led to a 3.7-fold increase (**Supplementary Fig. 3A**). Anti-CD40 plus IL-23 nominally increased IL-17 expression compared to IL-23 alone. In a similar assay using IL-17-GFP reporter B cells, adding IL-6, IL-21, or IL-17A to anti-IgM had minimal effect on IL-17A frequency compared to anti-IgM alone, and none synergized with anti-IgM plus IL-23 (data not shown).

Based on these findings, sort-purified TIM-4+ versus TIM-1^+^ B cells were stimulated with anti-IgM in separate cultures, and cytokine secretion was examined at 48h. TIM-4^+^ B cells stimulated with anti-IgM secreted IL-17A, but only with the addition of IL-23 (**Fig 3A**). In contrast, TIM-1^+^ B cells secreted minimal IL-17A even after adding IL-23. As a control for experiments described below, anti-IL-17A completely neutralized IL-17A detected in the supernatants. Next, we examined *Il17a* and *Il10* expression at the transcriptional level by performing quantitative PCR (qPCR) on sort-purified TIM-4**^+^**TIM-1**^-^**and TIM-1**^+^**TIM-4**^-^** B cells from WT or *Il23r^-/-^*mice after stimulation in separate cultures for 24 hours. Stimulation of WT TIM-4^+^ B cells with anti-IgM induced detectable *Il17a* mRNA, and this was increased ∼3-fold with the addition of IL-17A and ∼10-fold with IL-23 (**Fig. 3B**). Neutralizing IL-17 in the culture by addition of anti-IL-17A mAb, decreased *Il17a* mRNA expression induced by IL-23. This suggests that IL-17A secretion promotes its own expression in a positive feedback loop and that contributes to IL-23-mediated IL-17A transcription. TIM-4^+^ B cells from *Il23r^-/-^* mice had almost undetectable *Il17a* expression even after addition of exogenous IL-17, confirming the requirement for IL-23 signaling. Neither WT nor *Il23r^-/-^*TIM-1^+^ B cells expressed detectable *Il17a* mRNA.

**Fig 3.**
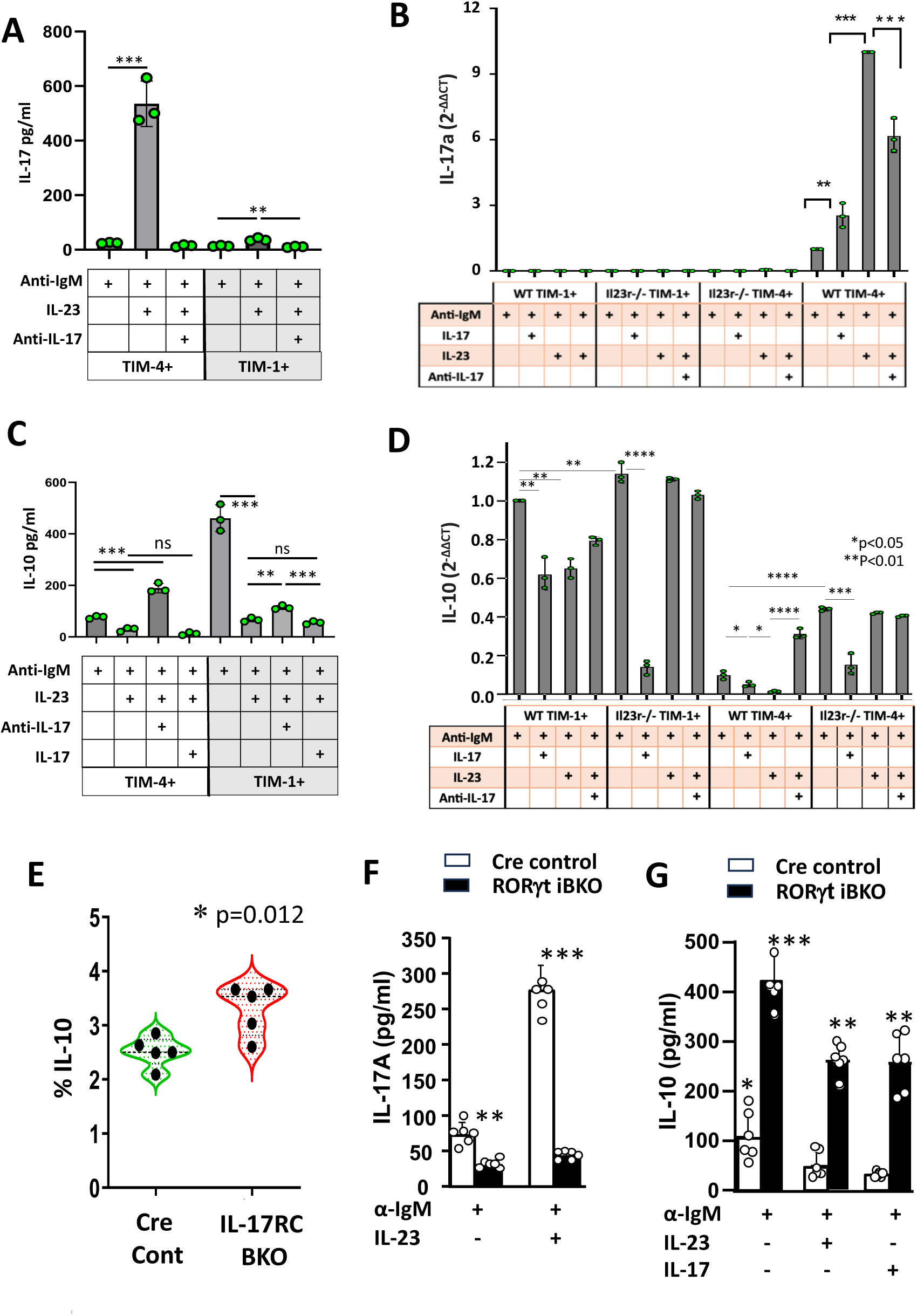
Regulation of IL-17A and IL-10 expression by IL-23 and IL-17. **(A)** TIM-1^+^TIM-4 and TIM-4^+^TIM-1^-^ CD19^+^ B cells from alloimmunized *Il23r^-/-^* and WT B6 mice were stimulated for 24 hours with anti-IgM and cytokines or neutralizing anti-IL-17A as indicated. IL-17A expression was compared to that of GAPDH by qPCR. The bar graph shows the relative expression of IL-17A mRNA as 2-^ΔΔ^CT performed in triplicate wells on cells pooled from 3 mice per group. **p<0.01; ***p<0.001. Data representative of three independent experiments. **(B-C)** TIM-1^+^TIM-4^-^ and TIM-4^+^TIM-1^-^ B cells from alloimmunized WT B6 mice were stimulated with anti-IgM alone or anti-IgM plus IL-23, IL-17A or anti-IL-17A neutralizing antibody as indicated. Bar graphs display the concentration of IL-17A (**B**) and IL-10 (**C**) in the supernatants (performed in triplicate wells on cells pooled from 3 mice per group). Data representative of three independent experiments. **p<0.01 and ***p<0.001. **(D)** TIM-1^+^TIM-4^-^ and TIM-4^+^TIM-1^-^B cells from *Il23r^-/-^* and WT B6 mice, were stimulated and IL-10 mRNA expression assessed by qPCR as in (**A**) above. The bar graph shows the relative expression of IL-10 as 2-^ΔΔ^CT in triplicate wells on cells pooled from 3 mice per group. **p<0.01; ***p<0.001. Data representative of three independent experiments. **(E)** Violin plot depicting mean frequency (+SD) of IL-10 expression (intracellular staining) on splenic CD19^+^ B cells from IL-17RC^fl/fl^CD19-Cre^+/-^ (IL-17RC-BKO) and CD19-Cre (Cre Control) mice. n=5 mice. *p<0.05. (**F, G**) Purified TIM-4^+^ CD19^+^ B cells from spleens of TAM-treated RORγT iBKO and Cre control mice were stimulated in vitro with anti-IgM alone, or anti-IgM plus IL-23 or IL-17A, for 48 hours. Bar graphs display the concentration of IL-17A (**F**), and IL-10 (**G**) in the supernatants, as measured by Cytometric Bead Array. * p <0.03 vs. other Control groups; ** p<0.001 vs. corresponding Control group; ***p<0.001 vs. all other groups; ****p<0.0001 vs. all other groups. (n=6 mice per group, in 2 independent experiments).

Next, we examined IL-10 secretion by sort-purified TIM-4^+^ versus TIM-1^+^ B cells. Anti-IgM-stimulated TIM-4^+^ B cells secreted only ∼1/6 as much IL-10 protein as TIM-1^+^ B cells (**Fig. 3C**), corresponding to the frequency of IL-10 expression determined by intracellular staining (**Fig. 1**). Although addition of IL-23 to anti-IgM promotes IL-17A secretion by TIM-4^+^ B cells, it inhibits their secretion of IL-10 by ∼3-fold (**Figs. 3A, C**). However, when IL-17A in the supernatant is neutralized with anti-IL-17A, anti-IgM plus IL-23-stimulated TIM-4^+^ B cells increase their IL-10 secretion almost 8-fold (reaching ∼2.5-fold higher than baseline with anti-IgM alone; **Fig. 3C**). Furthermore, IL-10 secretion after anti-IgM plus IL-23 treatment is further reduced by the addition of exogenous IL-17A. These data suggest that IL-17A directly inhibits IL-10 secretion by TIM-4^+^ B cells and plays a dominant role in the IL-10-inhibitory effect of IL-23.

Anti-IgM-treated TIM-1^+^ B cells secreted much more IL-10 than TIM-4^+^ B cells. Addition of IL-23 reduced IL-10 secretion by TIM-1^+^ B cells ∼8.5-fold (**Fig. 3C**). It is unlikely that IL-17A is the main mechanism by which IL-23 inhibits IL-10 secretion by TIM-1^+^ B cells, since these cells secrete minimal IL-17A (**Fig. 3A**), and unlike TIM-4^+^ B cells, neutralization of IL-17A only slightly restored IL-10 secretion by the TIM-1^+^ subset (**Fig. 3C**). Addition of IL-17A to IL-23 had no significant effect on IL-10 secretion compared to IL-23 alone (**Fig. 3C**).

Examination of *Il10* mRNA by qPCR generally corroborated cytokine secretion data (**Fig. 3D**). Neutralization of IL-17 in IL-23-treated WT B cells augmented *Il10* expression, particularly in the TIM-4^+^ subset which expresses more IL-17. *Il23r^-/-^* TIM-4^+^ and TIM-1^+^ B cells both expressed more *Il10* than their WT counterparts but did not respond to IL-17 neutralization, consistent with their lack of IL-17 expression (**Fig. 3B, D**). Yet, in the absence of exogenous/IL-23R signaling, addition of exogenous IL-17A reduced IL-10 expression in both TIM-4^+^ and TIM-1^+^ B subsets.

Taken together, these data indicate that inhibition of *Il10* expression by IL-23 is largely IL-17-dependent in TIM-4^+^ B cells and IL-17-independent in TIM-1^+^ B cells. However, IL-17A does directly inhibit *Il10* expression in both TIM-4^+^ and TIM-1^+^ B cells. If IL-17A directly inhibits B cell *Il10* expression, then IL-17R-deficient B cells should express more IL-10. To test this, we generated *Il17RC*^fl/fl^ X CD19-Cre (IL-17RC BKO) mice and compared these to CD19-Cre controls. Indeed, B cells from immunized IL-17RC BKO mice produced over 40% more IL-10 than Cre control B cells (**Fig. 3E**).

Finally, we addressed the degree to which regulation of cytokine secretion by IL-17 and IL-23 were RORγt dependent. Consistent with intracellular staining of freshly isolated cells (**Fig. 2G**), TIM-4^+^ B cells from TAM-treated RORγt iBKO mice secrete significantly less IL-17A in response to anti-IgM than those from Cre controls (**Fig 3F**). Moreover, *Rorc*-deletion essentially abrogated IL-17A expression induced by IL-23, confirming that IL-23 acts through RORγt to increase IL-17A in TIM-4+ B cells. The residual low-level IL-17A secretion in RORγt iBKO B cells is consistent with incomplete *Rorc* deletion (**Fig 2G**).

Consistent with intracellular staining (**Fig. 2I**), *Rorc* deficiency augmented IL-10 secretion by TIM-4+ B cells (**Fig. 3G**). Notably, exogenous IL-23 significantly reduced IL-10 secretion by *Rorc*-deficient TIM-4^+^ B cells indicating that in addition to IL-17-dependent inhibition of IL-10, IL-23 also inhibits IL-10 through RORγt- and IL-17- independent pathways (**Figs. 3D, G**). Exogenous IL-17 also inhibited IL-10 production independent of RORγt. However, even with the addition of exogenous IL-17 or IL-23, IL-10 secretion by RORγt-iBKO B cells remained far higher than in Cre-controls. This indicates that in addition to driving IL-17 expression, RORγt must inhibit IL-10 secretion through other (IL-17-independent) mechanisms. In this regard, we previously showed that RORγt can bind to the *Il10* promotor/enhancer elements and suppress IL-10 expression in Th17 cells (*22*).

### Profiling TIM-4^+^B cells reveals expression of multiple pro-inflammatory cytokines in a pattern resembling pathogenic Th17 cells

Given their opposing function and patterns of cytokine expression, we compared the transcriptomes of TIM-4+ and TIM-1+ B cells by population level RNA sequencing (RNA-seq). RNA was isolated from sort-purified TIM-4**^+^**TIM-1**^-^**, TIM-1**^+^**TIM-4**^-^**, and DN (TIM-4^-^TIM-1^-^) B cells from spleens of alloimmunized mice after 24-hour stimulation with anti-IgM plus IL-23. Principal component analysis (PCA) revealed that PC1 separated the expression profiles of TIM-1^+^ and TIM-4^+^ B cells, while PC2 separated DN B cells from both TIM-4^+^ and TIM-1^+^ B cells (**Supplementary Fig. 4A**). Variability in TIM-1^+^ samples from different mice along PC1 related to total number of unique genes detected. Comparison of TIM-4^+^ and TIM-1^+^ B cells by differential gene expression analysis revealed 3425 upregulated genes (613 with fold-change >1.5) and 7856 downregulated genes (7391 with fold-change >1.5) in the TIM-4^+^ population (FDR <0.05; **Fig. 4A)**. Genes of interest are displayed in the volcano plot (**Fig. 4B)**. Compared to TIM-1^+^ B cells, TIM-4^+^ B cells not only expressed higher levels of *Il17a*, but also a pro-inflammatory module comprising *Il17f, Il22, Csf2, Il6, Il1β, TNFα,* and *Il2* **(Fig. 4B-D**). As expected, TIM-1^+^ cells expressed more *Il10*, *Ctla4, Nte5* (expressing CD73), *ENTpd1* (expressing CD39), *Cd274* (expressing PD-L1), and *Pdcd1lg2* (expressing PD-L2)(*7*). *Ifng* expression showed a trend, but was not statistically higher in TIM-4^+^ B cells (**Fig. 4C**), even though IFNγ protein expression is higher in freshly isolated TIM-4^+^ versus TIM-1^+^ B cells(*9*). Surprisingly, in these stimulated cells, the TIM-4 gene (*Timd4*) was expressed at higher levels by TIM-1^+^ B cells even though TIM-4 protein was not detected by flow cytometry. Many of the same proinflammatory cytokines noted above were also increased in TIM-4^+^ B cells compared to DN or TIM-4^-^ B cells (**Fig. 4C, D**).

**Fig 4.**
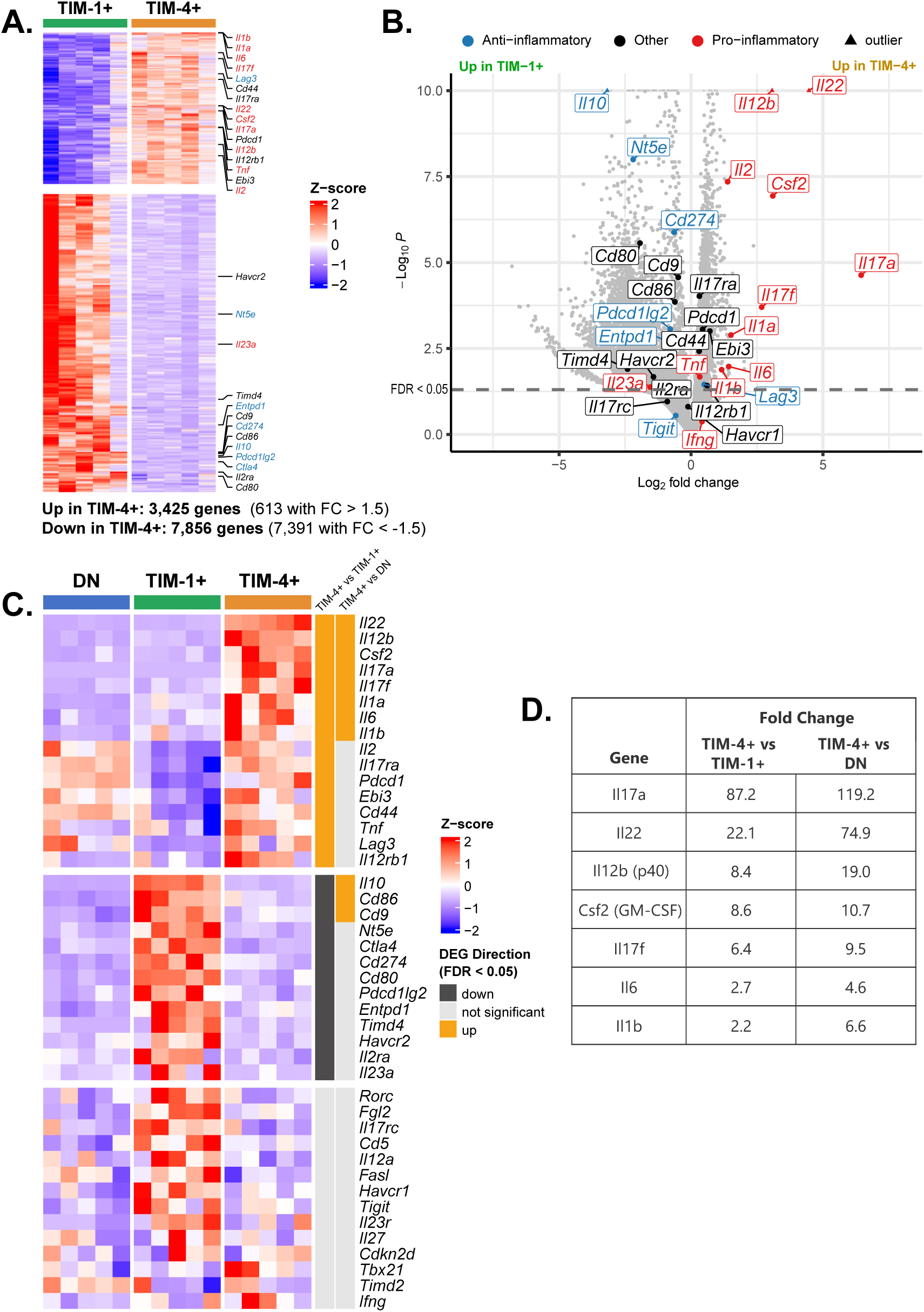
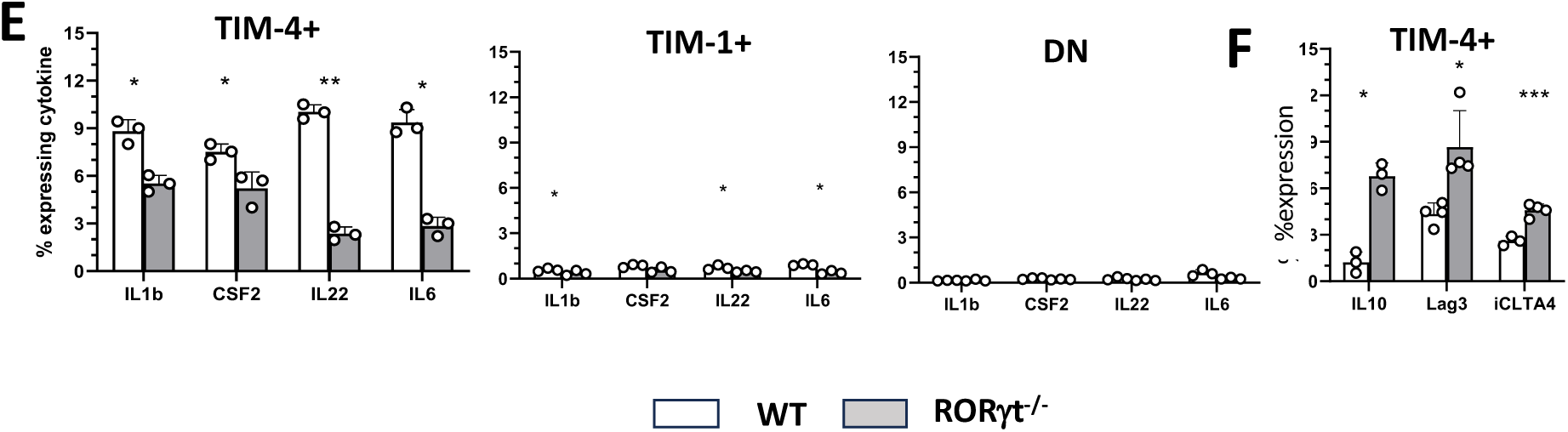
Differential gene expression in TIM-1^+^ and TIM-4^+^ B cells from alloimmunized mice. **A-B:** Amongst genes of interest, pro-inflammatory genes are depicted in red, anti-inflammatory genes are in blue, and other genes are in black text. **A**) Heatmap showing differentially expressed genes identified by RNA-seq (FDR<0.05). **B**) Volcano plot with selected genes of interest. The y-axis is capped at 25 for visual clarity. **C**) Heatmap comparing expression of selected genes of interest between three groups: TIM-1^+^, TIM-4^+^ and DN B cells annotated on the right. Vertical bars represent genes that are upregulated (orange), downregulated (black), or not statistically differentially expressed (gray, FDR ζ 0.05) in a given comparison between TIM-4^+^ B cells (T4) and either TIM-1^+^ (T1) or DN B cells. **D)** Fold change in expression between select genes in TIM-4^+^ vs. TIM-1^+^ B cells or TIM-4^+^ vs. all other (TIM-4^-^) B cells. Data are representative of a second independent experiment comparing TIM-4^+^ vs TIM-4^-^ B cells. (**E-F)** WT and RORγt KO mice were alloimmunized, and cytokine expression (IL-1β, CSF2, IL-22, IL-6, and IL-10) in TIM-1⁺, TIM-4⁺, and DN B cells, as well as LAG-3 and CTLA-4 expression in TIM-4⁺ B cells, was analyzed by flow cytometry. *p < 0.05; **p < 0.01; ***p < 0.001.

Intracellular staining confirms at the protein level, that TIM-4^+^ B cells express significantly more IL-1β, GM-CSF, IL-22, and IL-6 compared to TIM-1+ or DN B cells (**Fig 4E**). The production of each of these additional proinflammatory cytokines is partially RORγt-dependent. In addition to increased frequency of IL-10 production noted previously, *Rorc*-deletion increased expression of LAG-3 and CTLA-4 by TIM-4+ B cells (**Fig 4F**). These coinhibitory molecules are normally expressed by TIM-1+ Bregs(*7*). Other components of the TIM-1 regulatory module including CD73, TIGIT, CD39, PD1, PDL1, and PDL2, were not increased. Taken together, our data indicate that in addition to IFNγ, TIM-4^+^ Beffs express a RORγt-dependent proinflammatory module containing IL-17A and other proinflammatory cytokines resembling those expressed by pathogenic Th17 (pTh17) cells. The dominant regulatory function of *Rorc*-deficient B cells is likely due to the combined effect of decreased expression of multiple proinflammatory cytokines and an increase in expression of IL-10 and several other components of the TIM-1+ regulatory module.

Pathways analysis of the RNA-seq data utilizing the Gene set Ordinal Association Test (GOAT) revealed upregulation of multiple pathways in stimulated TIM-4^+^ versus TIM-1^+^ B cells that can be condensed into 4 domains: Immune activation; Stress/damage responses; Metabolic reprogramming; and Cell growth/proliferation and cell-cycle regulation (**Supplementary Fig 4B**). Specific enriched pathways included allograft rejection, TNF signaling via NFκB, and IFNα and IFNγ responses, consistent with heightened proinflammatory activity. For comparison, RNA-seq was also performed on freshly isolated (unstimulated) TIM-4+ and TIM-1+ B cells. Although the proinflammatory gene transcripts noted above were not upregulated in freshly isolated TIM-4+ B cells (data not shown), pathways analysis still identified enrichment of the allograft rejection and TNF signaling via NFκB pathways (**Supplementary Fig 4B**). However, six pathways related to activation signaling, proliferation, and metabolic reprogramming, that were upregulated in anti-IgM + IL-23-stimulated TIM-4+ B cells, were now downregulated. This suggests that TIM-4+ B cells are poised towards a stronger response to BCR-mediated activation signals than TIM-1+ B cells. The top 50 upregulated and downregulated genes in stimulated and unstimulated TIM-4^+^ versus TIM-1^+^ B cells are listed in **Table S1 and S2, respectively.**

Because TIM-4^+^ B cell cytokines closely resemble those expressed by pTh17 cells, we compared gene expression by TIM-4^+^ and TIM-1^+^ B cells to the pTh17 signature we previously identified in polarized pTh17 versus non-pTh17 cells (**Supplementary Fig. 4C)** (*22–24*). In addition to the cytokines pathognomonic for pTh17 cells like *Csf2* and *Il22*, TIM-4^+^ B cells exhibit increased expression of pTh17 signature genes including *Ccl3*, *Ccl4*, and *Ccl5*, *Stat4*, and *Lgals3* (encoding Galectin-3). However, other genes upregulated in this pTh17 signature, including, *Icos, Cxcl3, Gzmb, Casp1,* and *Tbx21* were not upregulated in TIM-4^+^ B cells. Amongst genes downregulated in pTh17 cells, *Il10, Maf, and Ikzf3* (IKAROS family zinc finger 3) are also downregulated in TIM-4^+^ B cells, while *Cd5l* was upregulated and *Ahr* expression was variable (**Supplementary Fig. 4D)**. Of genes identified in several other pTh17 signatures(*24–29*), only *Batf, Bhle40*, and *Cd44* were upregulated in TIM-4^+^ B cells, but not *Il23r*, the canonical Notch signaling mediator *Rbpj, IL1rn*, *Gpr65, Fcmr (Toso), Zbtb32 (Plzp), Batf3, Prdm1*, Protein receptor C (*Procr*), *Foxo1*, or *Irf4*, (**Fig 4C, Supplementary Fig. 4C**). These data suggest that while expressing similar cytokines, the regulation of TIM-4^+^ B cell induction and/or differentiation may be distinct from that of pTh17 cells.

## DISCUSSION

In addition to antibody production, B cells play an important role promoting or inhibiting the immune response (*1, 2, 4–6*). Our understanding of Breg biology was enhanced by the discovery of TIM-1 as a broad marker for Bregs that regulates their expression of an array of anti-inflammatory cytokines and coinhibitory molecules in addition to IL-10(*7, 8*). B cells producing various proinflammatory cytokines promote anti-microbial, tumor, allo-, and auto- immunity, yet only the phenotype of B cells expressing IFNγ has been reported (*2, 5, 6, 9, 12, 13, 30*). It was unclear whether B cells expressing different inflammatory cytokines shared a common phenotype or regulatory mechanisms. We now show that TIM-4 identifies proinflammatory Beffs that selectively express RORγt which underlies the expression of IL-17A as well as IL-17F, IL-22, IL-6, IL-1β, and GM-CSF. Thus, TIM-4^+^ is the first marker to broadly identify B cells expressing most of the proinflammatory cytokines previously attributed to B cell effector function and to our knowledge, this is the first description of RORγt expression by B cells. This allows us to begin to address how proinflammatory cytokine production is regulated in TIM-4^+^ Beffs and differs from the signals regulating TIM-1^+^ Breg activity.

It was previously reported that in the isolated setting of *T. cruzi* infection, a pathogen-specific transialidase alters CD45 glycosylation, and this induced RORγt-independent IL-17A expression in B cells(*14*). Whereas adoptive transfer of WT B cells into μMT mice enhanced T. cruzi clearance, adoptive transfer of *Il17*^-/-^ B cells had no effect. We now demonstrate that B cell IL-17A expression is a much more generalized component of B cell function, expressed in response to various types of immunization at a frequency similar to that of B cell-derived IL-10. In these settings, IL-17A is RORγt-dependent and selectively expressed by TIM-4^+^ B cells where it regulates expression of a pTh17-like pattern of cytokines. In addition to IL-23-induced proinflammatory cytokines, the TIM-4^+^ pro-inflammatory module shares additional similarities with pTh17 cells including chemokines (*Ccl3, Ccl4, Ccl5), Lgals3* (encoding Galectin3), and *Stat4* (*23*). Despite this resemblance, the regulation of proinflammatory cytokine versus IL-10 expression by TIM-4^+^ B cells, and between TIM-4^+^ Beffs and the opposing TIM-1^+^ Breg subset, differs from that seen in non-pathogenic Th17 (npTh17) versus pTh17 cells. npTh17 cells express IL-10 along with IL-17A(*31, 32*). Although npTh17 cells are not frankly suppressive, they play an essential role maintaining the integrity of the gut mucosal barrier. In contrast, pTh17 cells lack IL-10, but express IL-17A in conjunction with a host of other pro-inflammatory cytokines including GM-CSF which plays an important inflammatory role in EAE (*32, 33*). The equilibrium between npTh17 and pTh17 cells is carefully regulated by balancing IL-10 and IL-23R expression, respectively. For example, compared to pTh17 cells, npTh17 cells express increased CD5L, which interacts with fatty acid synthase to alter lipid biosynthesis and reduce RORγt-agonist ligands(*34*). This reduces RORγt-DNA binding, resulting in decreased activation of the *Il23r* locus and decreased repression of the *Il10* locus, which together, favor the differentiation of IL-10-expressing npTh17 rather than pTh17 cells. In contrast, proinflammatory TIM-4^+^ Beffs (which express little *Il10*), express more *Cd5l* than TIM-1^+^ Bregs. It is possible that CD5L acts to modulate TIM-4^+^ Beff proinflammatory activity. As a further distinction, in Th17 cells, increased RBPJ (a mediator of NOTCH signaling), enhances *Il23r* expression which drives pTh17 differentiation (*35*). Yet, neither *Rbpj* nor *Il23r* are differentially expressed by TIM-4^+^ versus TIM-1^+^ B cells, which is consistent with the IL-23 responsiveness of both subsets. Indeed, IL-23R signaling reduces IL-10 expression in both TIM-1^+^ and TIM-4^+^ B cells. Yet, IL-23 only induces inflammatory cytokine expression in TIM-4^+^ B cells, likely due to their almost exclusive expression of RORγt. In addition to its requisite role in IL-17A expression, RORγt augments the expression of other proinflammatory cytokines by TIM-4^+^ Beffs, as in pTh17 cells. TIM-4^+^ B cells also differentially express *Bhle40*, a basic helix-loop-helix transcription factor that promotes *Csf2* and inhibits *Il10* expression, and *Batf*, an AP-1 transcription factor that transactivates *Il17* and *Il22* expression in pTh17 cells(*36, 37*). Thus, TIM-1^+^ and TIM-4^+^ B cells with opposing functions as Beffs and Bregs, respectively, express transcription factors that differentially regulate regulatory and proinflammatory cytokine loci.

Our analysis revealed that IL-17A directly inhibits *in vitro* IL-10 expression by both TIM-4^+^ and TIM-1^+^ B cells (**Fig. 3D, G**). To our knowledge, a pathway leading from IL-17A signaling to inhibition of IL-10 expression has not been previously described. While exogenous IL-17 inhibits B cell-expressed IL-10 in vitro, B cells from mice that specifically lack B cell-derived *Il17* or *Il17rc* express significantly more IL-10 (Figs. 2, 3E). This indicates that *in vivo,* IL-17 plays an autocrine role inhibiting B cell IL-10 production and suggests that TIM-4^+^ B cells must largely reside near other B cells and away from other endogenous sources of IL-17. IL-23 also strongly inhibits IL-10 production by TIM-4^+^ B cells, even in the absence of RORγt, indicating an IL-17-independent effect (Fig. 4D). However, in WT TIM-4^+^ B cells, IL-10 expression is markedly enhanced by neutralizing IL-17, even in the presence of exogenous IL-23 and intact RORγt (Fig. 3C, D). This suggests that IL-17 plays a dominant role maintaining the suppressing of IL-10 in TIM-4^+^ B cells. The situation is somewhat different in TIM-1^+^ B cells. While both IL-17A and IL-23 can independently inhibit IL-10 expression by TIM-1^+^ B cells (Fig. 3D), here, the IL-17 pathway does not predominate. IL-23 strongly inhibits IL-10 expression by TIM-1^+^ B cells despite barely detectable IL-17A expression, and IL-17A neutralization or addition of exogenous IL-17 have little effect. Finally, in addition to its requisite role in IL-17A expression, RORγt plays an IL-17 and IL-23 independent role inhibiting IL-10 expression in TIM-4^+^ B cells (Fig 4D). This is consistent with our previous findings that RORγt can bind to the IL-10 promotor /enhancer elements and inhibit its expression (*34*).

In summary, in TIM-4^+^ B cells, IL-23 drives the expression of a proinflammatory module which includes RORγt-dependent IL-17A production. IL-17A acts in an autocrine manner to enhance its own expression, and along with RORγt, suppresses IL-10 and enforces pro-inflammatory Beff activity. In contrast, IL-23 has minimal effect on proinflammatory cytokine expression by TIM-1^+^ B cells which lack RORγt, while IL-23 and IL-17A independently inhibit IL-10 expression. These findings suggest that cross-regulation of Bregs and Beffs may occur at several levels. IL-23 in the microenvironment should inhibit *Il10* expression by TIM-1^+^ B cells and augment expression of the TIM-4^+^ proinflammatory module. IL-17A expressed by TIM-4^+^ B cells not only enforces their Beff activity but may help suppress *Il10* expression by TIM-1^+^ Bregs. Finally, impaired IL-17 production by TIM-4^+^ Beffs should increase their *Il1*0 expression and Breg activity. Thus, we have identified a reciprocal relationship between B cells with regulatory and effector activity. In this regard, we recently showed that deletion of TIM-1 expression in B cells inhibits tumor progression and results in the expansion of B cells in the draining LNs that express proinflammatory cytokines in a type I IFN-mediated manner(*38*). Whether this is due to the acquisition of inflammatory function by B cells that were destined to become TIM-1^+^ Bregs, or due to increased TIM-4^+^ Beffs in the absence of TIM-1^+^ Bregs, is not yet clear. In conclusion, TIM-4 is a broad marker for RORγt-expressing B cells that exhibit Beff function. IL-23 and IL-17A selectively enhance proinflammatory cytokine expression by TIM-4^+^ B cells, while preventing their conversion into IL-10-expressing B cells with regulatory activity. Further elucidation of these signals may allow therapeutic manipulation of TIM-4^+^ B cells to appropriately inhibit or enhance the immune response in settings of autoimmunity, transplantation, and cancer.

## METHODS

### Study Design

To further study the role of TIM-4^+^ B cells, we examined their cytokine production by flow cytometry and confirmed results using IL-17-reporter mice, and in vitro cytokine secretion. The role of B cell IL-17 was studied in EAE and islet allograft models by B cell adoptive transfer, generation of BM chimeras, and inducible B cell specific RORγt knockouts. These studies revealed an inverse relationship between IL-17 and IL-10 expression. The role of various cytokines in regulating IL-17 vs. IL-10 expression was directly tested in Tim-4^+^ as well as Tim-1^+^ Bregs in vitro by RT-PCR and cytokine secretion assays. Finally, Bulk RNAseq was performed to reveal that Tim-4^+^ Beffs express a module of proinflammatory cytokines, allowing a comparison to the genes signature of pTh17 cells. Previously published methods and those not used in the main Figs. are described in Supplementary Methods.

### Mice

C57BL/6 (B6; H-2b), BALB/c (H-2d), *Rorc*^-/-^, *Rorc^fl/fl^*^l^, *Il17a^−/−^*, B6 IL-17A–EGFP reporter, OT-I transgenic, OT-II transgenic and μMT (B6) mice were from The Jackson Laboratory. *hCd20-ERT2.Cre* (from Mark Shlomchik, University of Pittsburgh(*39*) were crossed with Rorc^fl/fl^ mice to generate *Rorc^fl/fl^*.*hCd20Cre*^+/-^ (RORγt iBKO) mice. Act-mOVA F1 mice (from Fadi Lakkis, University of Pittsburgh) were generated by breeding BALB/c with B6.*Act-OVA* mice(*20*). Mice were used at 6–10 weeks of age and were housed with food and water ad libitum.

### Islet transplantation and alloantigen immunization

400 allogeneic islets from BALB/c donors were transplanted under the left renal capsule of sex-matched B6 recipients with streptozotocin-induced diabetes, as previously described (See Supplementary Methods)(*8, 9*). In some experiments, mice were alloimmunized by i.p. injection of 2 × 10^7^ mitomycin C–treated BALB/c splenocytes(*8, 9*).

### Flow cytometry

Fluorochrome-conjugated mAbs were purchased from BD Biosciences, eBioscience, and BioLegend. Staining was performed in the presence of Fc block(anti-CD16/CD32) and LIVE/DEAD Fixable Aqua viability indicator as described (*8, 9*). Flow acquisition was performed on LSRII analyzers (BD Biosciences), and data were analyzed using FlowJo software (BD). Background staining was determined with isotype-matched controls or GFP^-^littermates. Detection of intracellular cytokine detection by flow cytometry was performed as previously described (*8, 9*). Briefly, T cells were cultured 4 h with PMA (50 ng/ml), ionomycin (500 ng/ml), and GolgiPlug (1 μl/ml). For IL-10, B cells were cultured 5 hours with LPIM (LPS (10 μg/ml), PMA, ionomycin, and monensin (1 μl/ml)). For IL-17, and other proinflammatory cytokines, B cells were cultured 5 hours with PIB (PMA, ionomycin, and Brefeldin A (1 μl/ml)). Cells were permeabilized using intracellular staining kits from BD Biosciences or eBioscience.

### Cell preparation and adoptive transfer

B cells were isolated from spleens of naïve or alloimmunized (d14) WT or IL17^-/-^ mice by negative selection (EasySep; StemCell Technologies; purity >98%). CD19^+^TIM-4^+^ and CD19^+^ TIM-4^−^ B cells were subsequently isolated by FACS using BD FACSAria (>95% purity). 5–7 ×10^6^ purified B cell subsets from naïve or alloimmunized syngeneic mice were adoptively transferred (i.v.) into otherwise untreated µMT recipients followed by islet transplants or EAE induction.

### EAE

Mice were immunized s.c. with MOG 35–55 (200µg) and CFA (200 µl; Difco Laboratories) and received pertussis toxin (200 ng/mouse; i.p.) on days 0 and 2 (*7*). Mice were scored daily for severity of EAE, as described (See Supplementary Methods) (*7*).

### Bone marrow (BM) chimeric mice

μMT (H-2b) mice were lethally irradiated (1000 rad) and reconstituted with 80:20 mixtures of syngeneic BM cells from: μMT plus WT, μMT plus *Il17a^−/−^*, or μMT plus *Rorc^−/−^* donors (*9*). BM chimeras were used after 8 weeks to allow immune reconstitution. B cells were fully reconstituted (from non-μMT BM).

### Cytokine Secretion

IL-17A and IL-10 concentration in supernatants of B cells/subsets was determined by Mouse Cytokine Cytometric Bead Array (CBA) per manufacturer’s instructions (BD Biosciences) after 48 hours of stimulation with anti-IgM plus various cytokines. PMA and ionomycin were added for the final 5h. 3000 events were acquired per sample on an LSR II (BD) analyzer.

### RNA isolation, real-time PCR

Splenic CD19^+^ B cells, isolated from alloimmunized B6 mice using the EasySep™ Mouse Pan-B Cell Isolation Kit (STEMCELL; purity >98%), were subsequently sorted based on staining criteria as CD19^+^ TIM-1^+^TIM-4-, TIM-4^+^TIM-1-, and DN (DUMP-negative for CD3, CD64, Gr-1, and Ter119) populations. These sorted B cells were then stimulated with anti-IgM along with various cytokines for 24 hours, with the addition of PMA and ionomycin during the final 3 hours of stimulation. Total RNA was extracted using RNeasy Micro Kit (Qiagen). 100ng of total RNA was used for cDNA synthesis by RNA to cDNA EcoDry™ Premix (Double Primed) kit (Takara), and qPCR was performed using TaqMan primers with TaqMan™ Fast Advanced Master Mix on StepOnePlus Real-Time PCR Systems (Applied Biosystems). Relative expression was calculated using the ΔΔCt method and level was normalized to *GAPDH*. TaqMan primers and probes were from Applied Biosystems: Mm00439618_m1 (*Il7A*), Mm01288386_m1(*Il10*), Hs02786624_g1(*Gapdh*).

### RNA-seq

B cells were purified and stimulated as above except sorting was repeated 2-3X until the purity was ∼100%. Total RNA quality was checked using a Bioanalyzer-2100 (Agilent Technologies) and submitted to the University of Pittsburgh Health Sciences Sequencing Core. Libraries were prepared (Ultra-low input RNA-seq Preparation Kit; Takara) and subjected to paired end sequencing (2 × 75 bp) using Illumina/NextSeq. Data was returned as NextSeq Fastq files.

### Analysis of bulk RNA-seq data

Bulk RNA-seq data were preprocessed on the Google Cloud Platform via the Terra scientific computing platform (Terra.Bio). Briefly, raw reads were aligned to the mouse genome mm10 using the STAR (v2.7.5a) alignment tool. Duplicate reads were identified using Picard MarkDuplicates. RNA-SeQC 2 was used for assessing sequencing depth and mapping quality. Raw counts and transcript per million (TPM) estimates were quantified using RSEM. All samples passed quality control and were included in subsequent analyses.

Downstream analyses were carried out in R (v4.1.3). Principal Component Analysis (PCA) was conducted using the R function *prcomp*. The gene with the highest PC2 loading, *AY036118*, was removed since it represents ribosomal RNA contamination(*40*). Differential gene expression analysis was performed using DESeq2 (v1.32.0). Lowly expressed genes with a total count lower than 10 were prefiltered. P-values were adjusted by Benjamini-Hochberg (BH) method(*41*), and log2 fold changes were shrunk using the ‘apeglm’ algorithm(*42*). Shrunken fold changes are depicted in all Figs. Heatmaps were created using R package ComplexHeatmap (v2.13.1) on normalized counts (normalized using the *counts* function from the BiocGenerics package (v0.40.0)(*43*) and were scaled by row. Volcano plots were generated using EnhancedVolcano (v1.24.0).

### Statistics

Statistical analyses of allograft survival used log-rank (Mantel-Cox) test. Differences were considered to be significant at *P* values <0.05. The clinical score and incidence of EAE were analyzed by 2-way Anova, and comparisons for results (mean ± SEM) for cytometric bead array. FACS, and real-time PCR were analyzed by Student’s 2-tailed *t* test with p <0.05 considered significant. Paired 2-tailed t-test was used when cells from the same mouse were compared. Significantly differentially expressed genes for RNA-seq analysis were identified using DESeq2 with Benjamini-Hochberg adjusted p-value <0.05. No statistical methods were used to predetermine sample sizes, but our sample sizes are standard for the field(*7, 8*). No data were excluded unless stated otherwise. Data distribution was assumed to be normal, but this was not formally tested.

### Study approval

Animal studies were approved by the Institutional Animal Care and Use Committee at the University of Pittsburgh (Pittsburgh, PA, USA).

## Data availability

All bulk RNA-seq data have been deposited in the NCBI’s Gene Expression Omnibus (GEO) database and are publicly available under accession number GSE253925. All raw data and materials will be made available to investigators upon request.

## Acknowledgements

We would like to thank Dr. Mark Shlomchik for supplying the hCD20.ERT2.Cre mice. Funding: This research was supported by NIH grants R01 AI114587 and AI184980.

## Author contributions

QD and DMR conceived the project. QD designed and performed most biological experiments. DMR supervised the study with key input from VKK. JLG provided data. AZ, ET, and YW performed, and AS supervised, computational analysis. QD and DMR analyzed the data. QD, JLG, VKK, and DMR wrote the paper with input from all the authors.

aaa

---

**Supplementary Fig.1:**
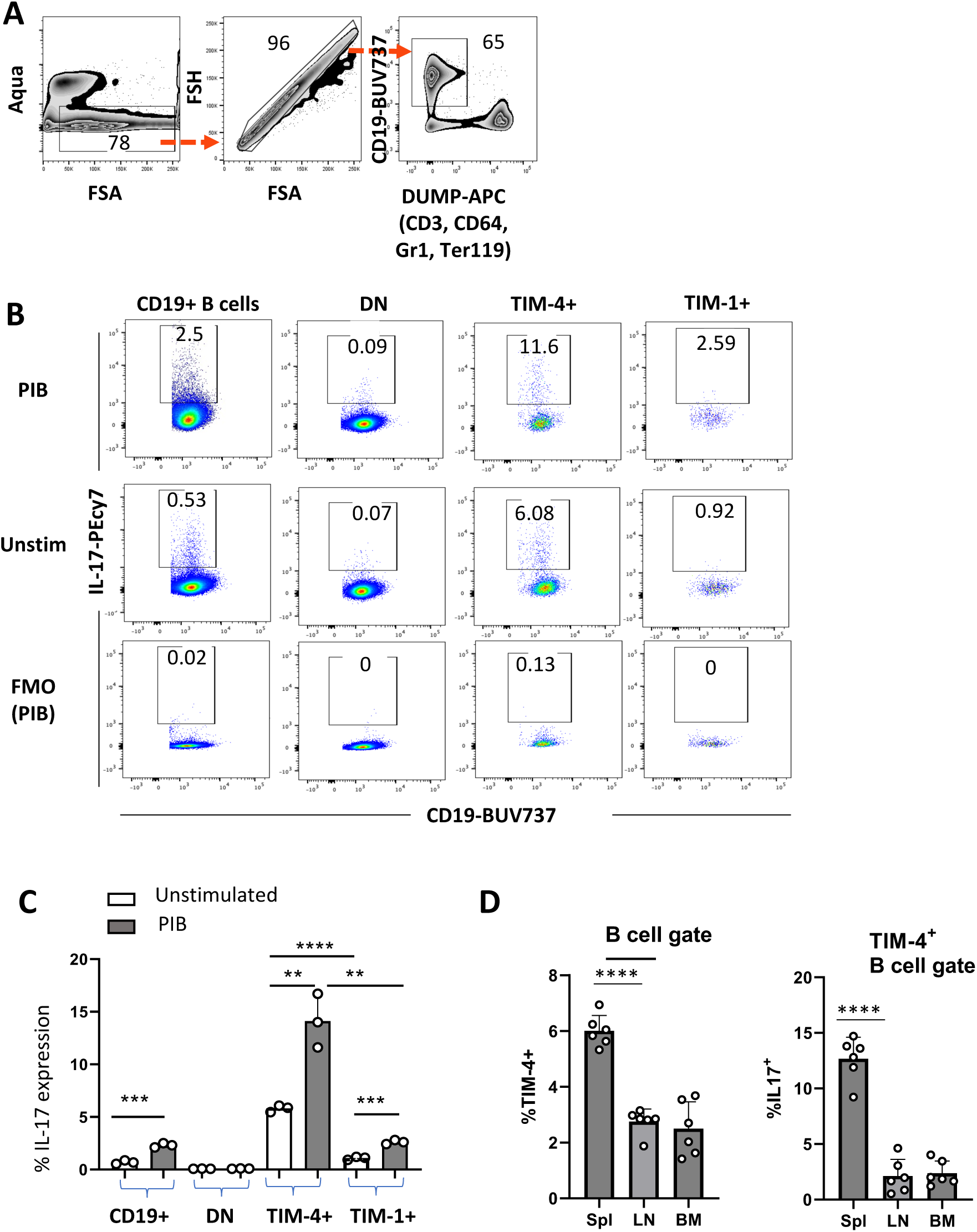
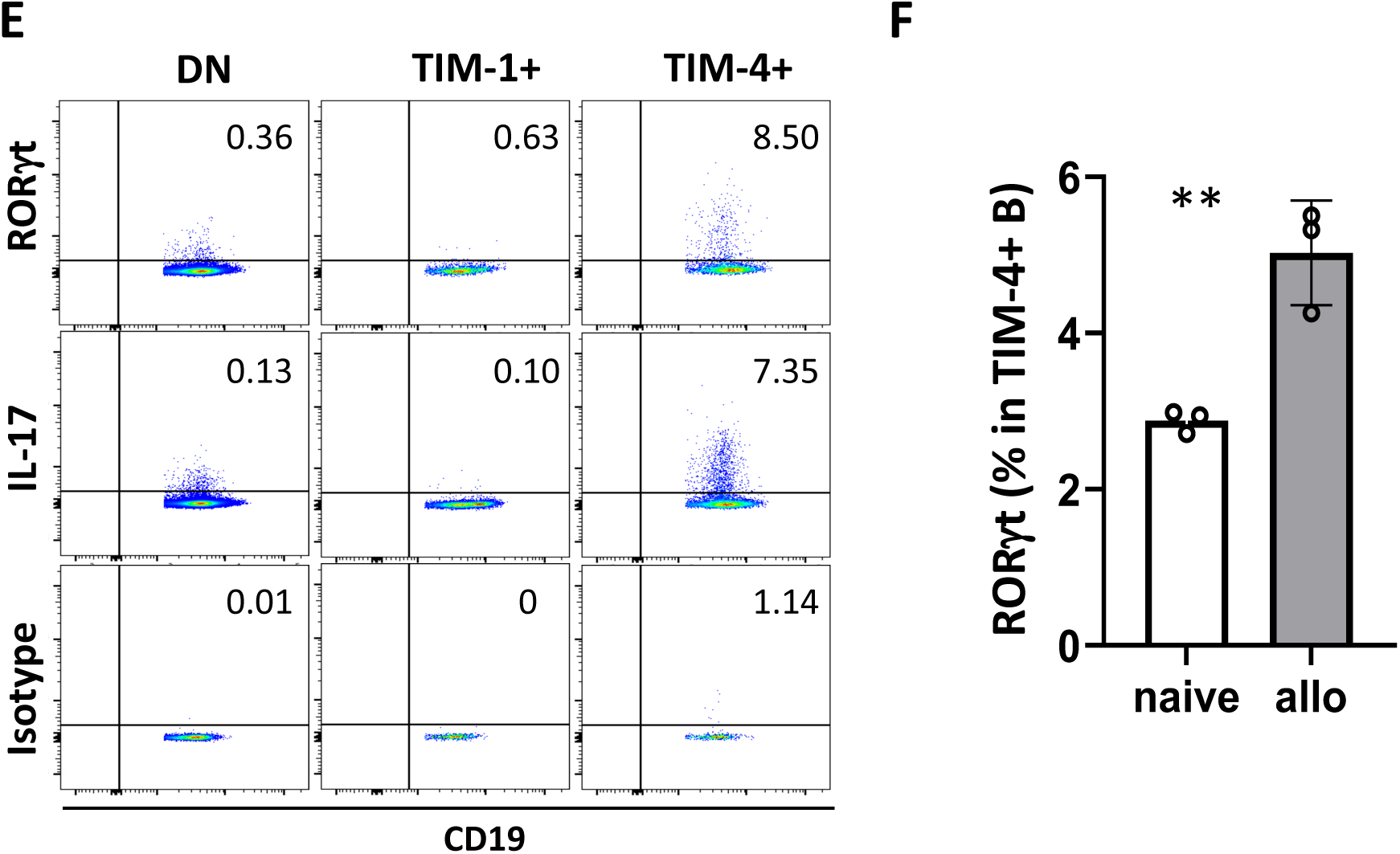
Preferential IL-17A expression on TIM-4^+^ B cells. **(A)** Representative FACS plots for splenocytes from alloimmunized Wt B6 mice. Gating strategy for selection of CD19^+^ Dump-gate (CD3, CD64, Gr1, Ter119) negative B cells assessed for IL-17A and TIM-4 expression in Figure 1E. (**B**) Representative FACS plots of IL-17A expression on total CD19^+^ B cells, and TIM-1^-^TIM-4^-^ (DN), TIM-1^+^, and TIM-4^+^ B cell subpopulations from spleens from mice 14 days after immunization with MOG_35-55_, CFA, and pertussis toxin to induce EAE (unstimulated vs. PIB for 5 hours). **(C)** Bar graph showing mean frequency (+ SD) of IL-17A expression by the B cell subpopulations shown in **(B)** (n≥3 mice per group). (**D**) Bar graphs showing the mean frequency (± SD) of TIM-4 expression in CD19⁺ B cells and IL-17A expression in TIM-4⁺ B cells from the spleen, lymph nodes, and bone marrow of alloimmunized WT B6 mice. (n=3 mice per group, in 2 independent experiments). **p <0.01, ***p<0.001, ****p<0.0001 versus indicated groups. (**E-F**) RORγt is selectively expressed by TIM-4^+^ B cells, and this correlates directly with IL-17A expression. Splenic B cells from C57BL/6 mice were assessed 14 days after alloimmunization. (**E**) Representative flow cytometry plots show RORγt and IL-17A expression on total DN (TIM-1^-^TIM-4^-^), TIM-1^+^, and TIM-4^+^ B cell subpopulations. isotype (control) shown in bottom row. (**F**) Bar graph showing the mean frequency (± SD) of RORγt expression in the TIM-4⁺ B-cell subpopulation from naïve and alloimmunized B6 mice.

**Supplementary Fig. 2:**
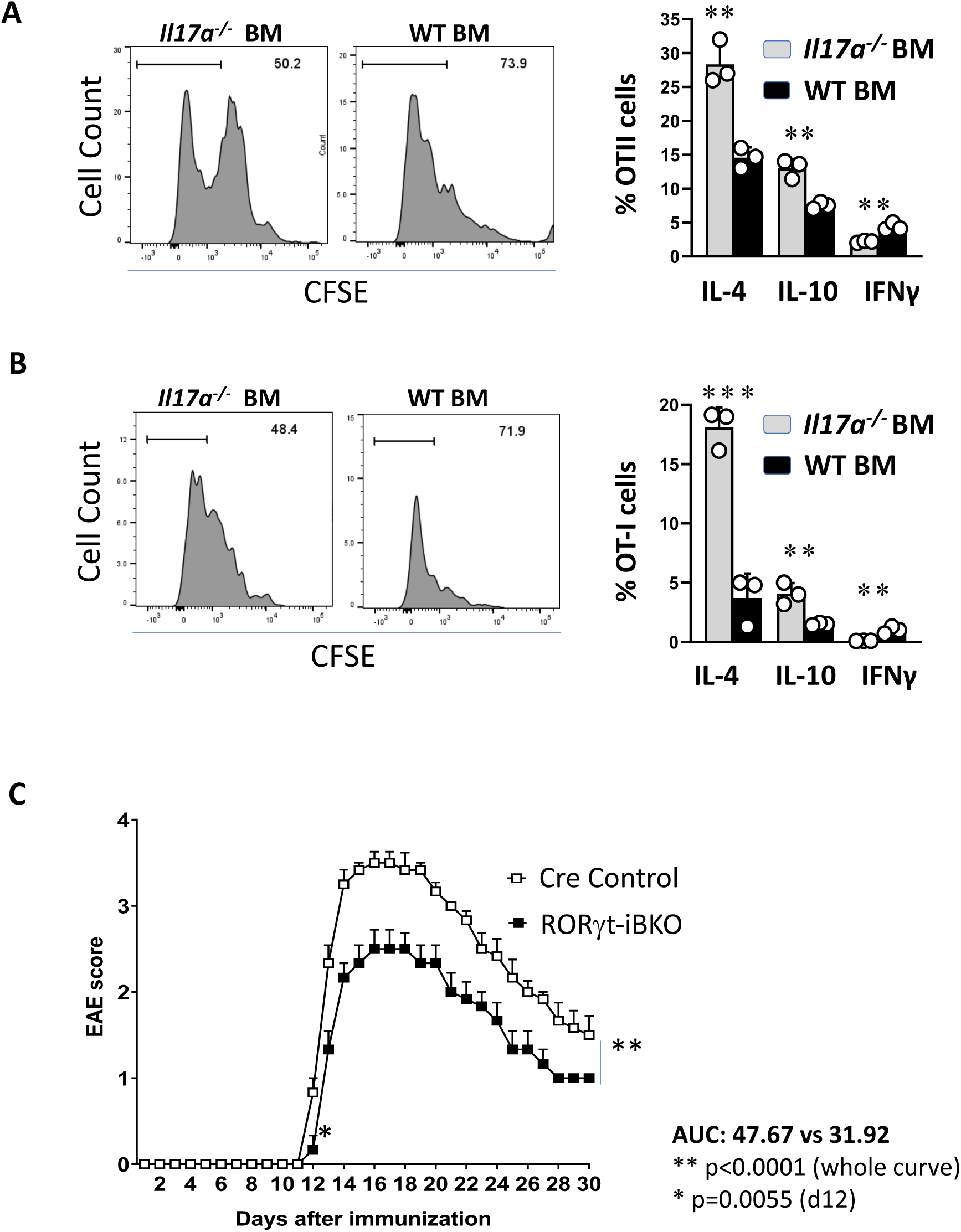
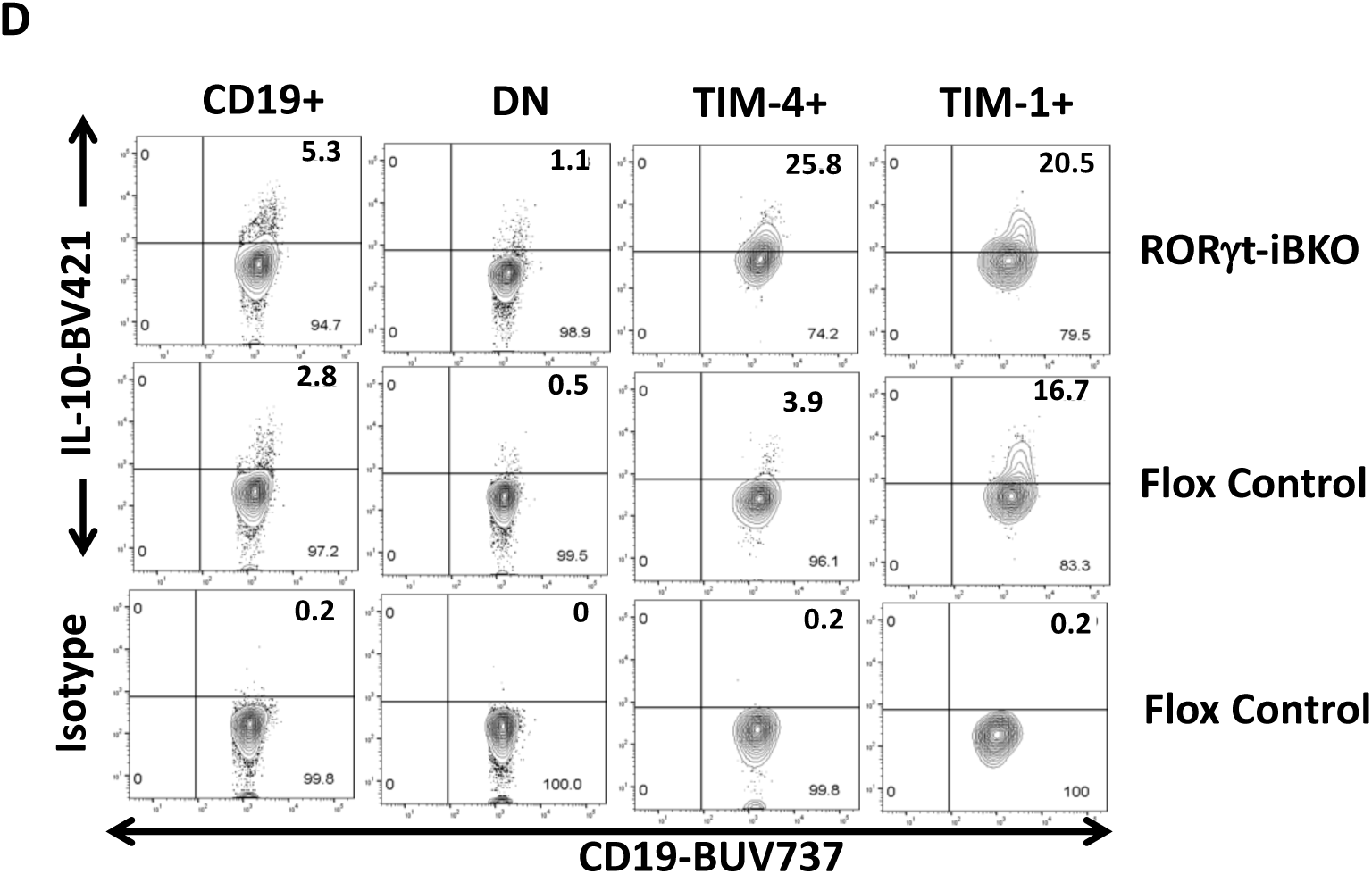
The Role of B cell IL-17 and RORγt on the immune response. **A and B:** BM chimeras were generated as in **Fig 2F**, by reconstituting lethally irradiated μMT mice with either 80% μMT plus 20% either *Il17a^−/−^* or WT bone marrow. After 8 weeks, BM chimeras were immunized with mitomycin C-treated splenocytes from Act-OVA.BALB/cXB6 F1 mice followed by adoptive transfer of 2×10^6^ CFSE-labeled CD90.1^+^ OT-II CD4^+^ **(A)** or CD90.1^+^ OT-I CD8^+^ T cells **(B)**. Transferred (CD90.1^+^) OT cells in spleen from the recipient mice were analyzed by flow cytometry for proliferation by CFSE dilution (day 4; Left panels), and intracellular cytokines expression (day 7; Right panels). n=3 mice per group. *p <0.05, **p <0.01 **(C)** Tamoxifen-treated RORγt-iBKO and Cre control mice were immunized to induce EAE and monitored daily for clinical signs (n=8-10 mice/group). Differences in clinical score was assessed using 2-way ANOVA (****p < 0.0001). **(D)** Representative flow cytometry plots show IL-10 expression (Intracellular staining after LPIM stimulation) on total CD19^+^ B cells, and TIM-1^-^TIM-4^-^DN, TIM-1^+^ and TIM-4^+^ B cell subpopulations from spleens of tamoxifen-treated alloimmunized RORγt-iBKO and Flox control mice.

**Supplementary Fig. 3:**
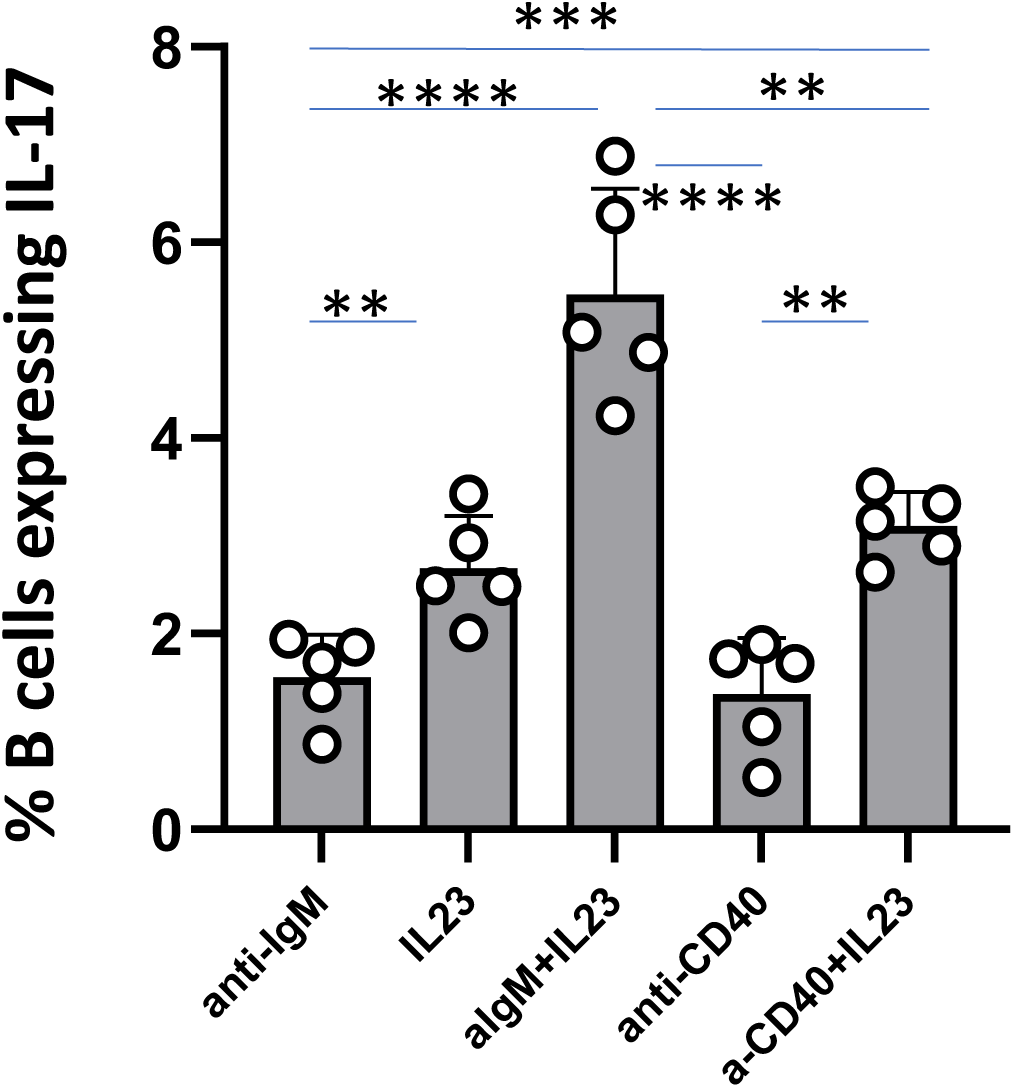
B cell IL-17A expression i*n vitro* is increased by addition of IL-23. Splenocytes from WT B6 mice were stimulated with anti-IgM or anti-CD40 in the presence or absence of IL-23 for 24 h, and B-cell IL-17A expression was assessed by intracellular flow cytometry (CD19⁺ gate). Statistical significance: *p<0.05; ** p<0.01; *** p<0.001; **** p<0.0001.

**Supplementary Fig.4:**
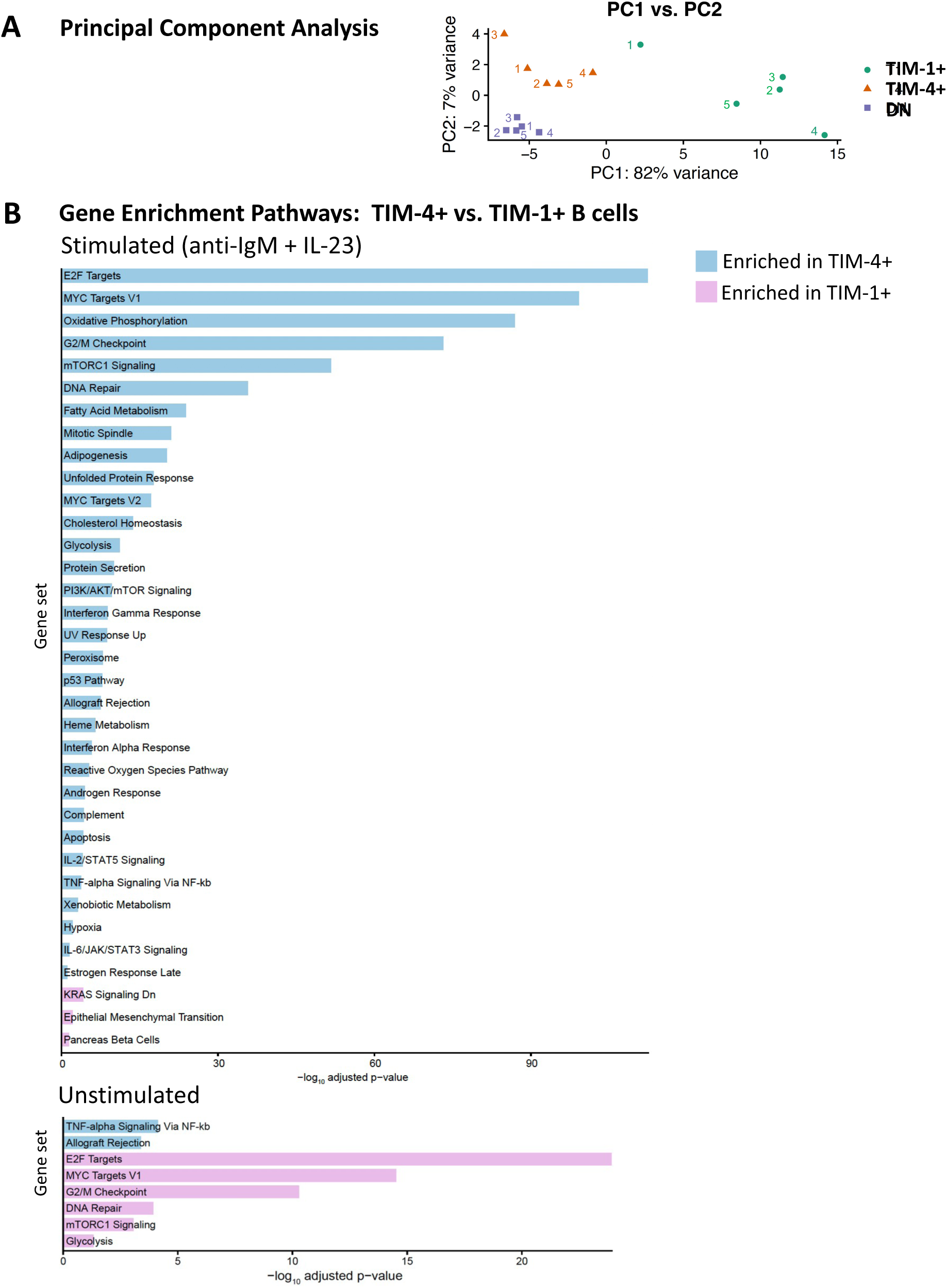
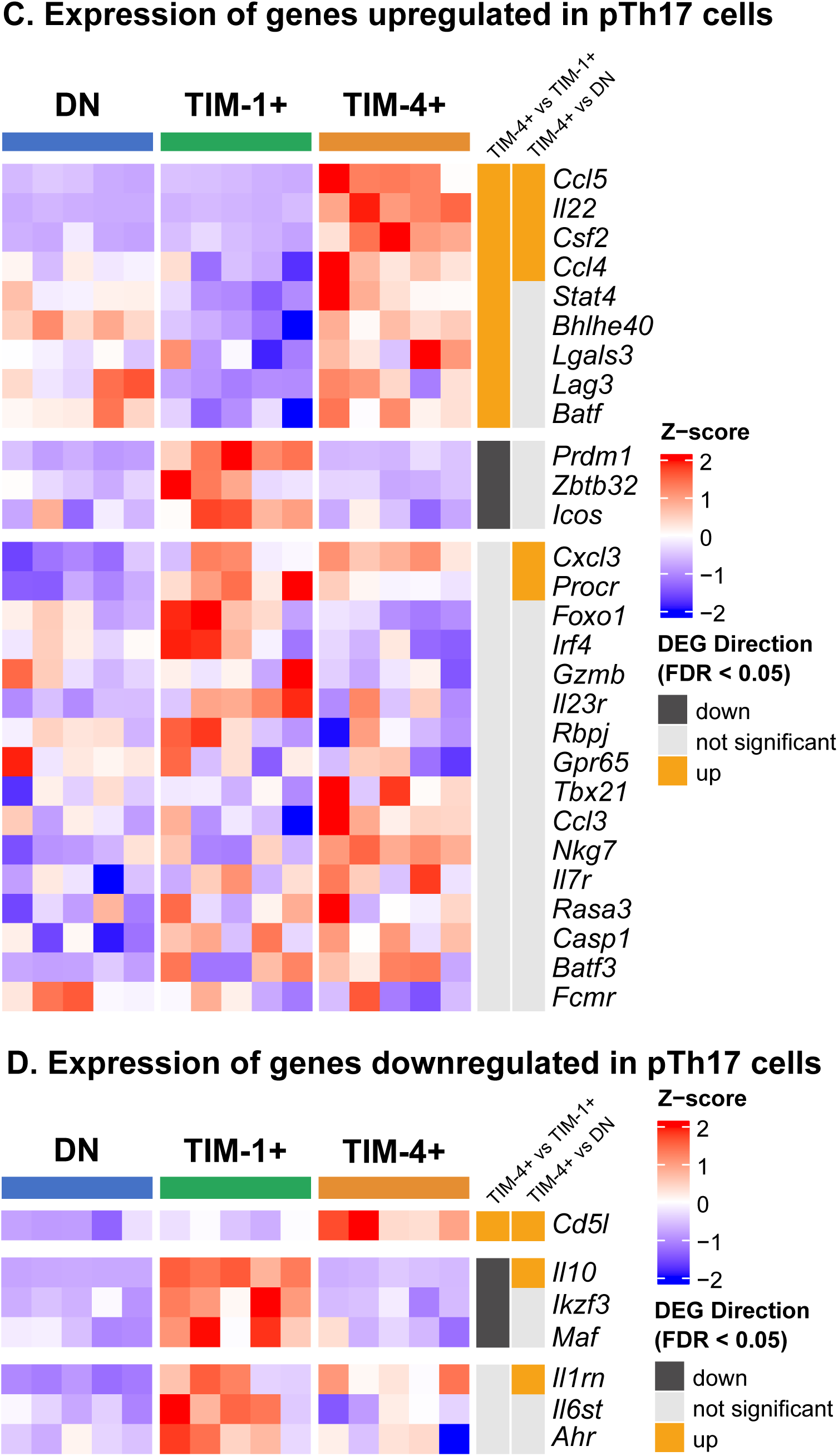
Gene expression in TIM-1^+^, TIM-4^+^, and DN B cells. **(A)** Principal Component Analysis (PCA) reveals TIM-1^+^, TIM-4^+^, and DN B cells comprise 3 distinct cell populations. PCA plot derived from bulk RNA-seq showing the relationship between the gene expression profiles of TIM-1^+^TIM-4^-^CD19^+^, TIM-4^+^TIM-1^-^CD19^+^, and TIM-1^-^TIM-4^-^CD19^+^ (DN) B cells from day 14 alloimmunized mice stimulated with anti-IgM plus IL-23 (24 hours) and PMA and ionomycin (3 hours). Each point represents a sample from an individual mouse, labelled (1–5) according to corresponding mouse. **(B)** Gene set Ordinal Association Test (GOAT) was used to identify gene set enrichment for differentially expressed genes in B cells stimulated with anti-IgM and IL-23. Enriched Hallmark pathways are ranked by –log₁₀ adjusted *P* value. All Hallmark gene sets with FDR ≤ 0.05 are shown. Stimulation induced strong enrichment of MYC and E2F target genes, G2/M checkpoint, oxidative phosphorylation, and mTORC1 signaling, consistent with IL-23–driven metabolic activation and proliferative programming. GOAT of unstimulated B cells showing enrichment of inflammatory and homeostatic pathways, including TNF-α signaling via NF-κB and allograft rejection signatures, alongside limited activation of cell-cycle–associated gene sets. These data suggest that IL-23 stimulation shifts B cells from a basal inflammatory state toward a metabolic and proliferative transcriptional program. **(C,D)** Heatmaps showing comparison of gene expression assessed by RNA-seq for genes identified as being upregulated (**C**) or downregulated (**D**) in pTh17 cells (pTh17 signature) in various publications (*23, 25–29*). Genes that are differentially expressed (FDR <0.05) between TIM-4^+^ B cells and either TIM-1^+^ B cells or DN B cells are color coded (orange for up-regulated and black for down-regulated; gray if not statistically significant) by corresponding vertical lines at the right of each heatmap. The gene expression values are row normalized (z-score).

**Supplementary Table 1:**
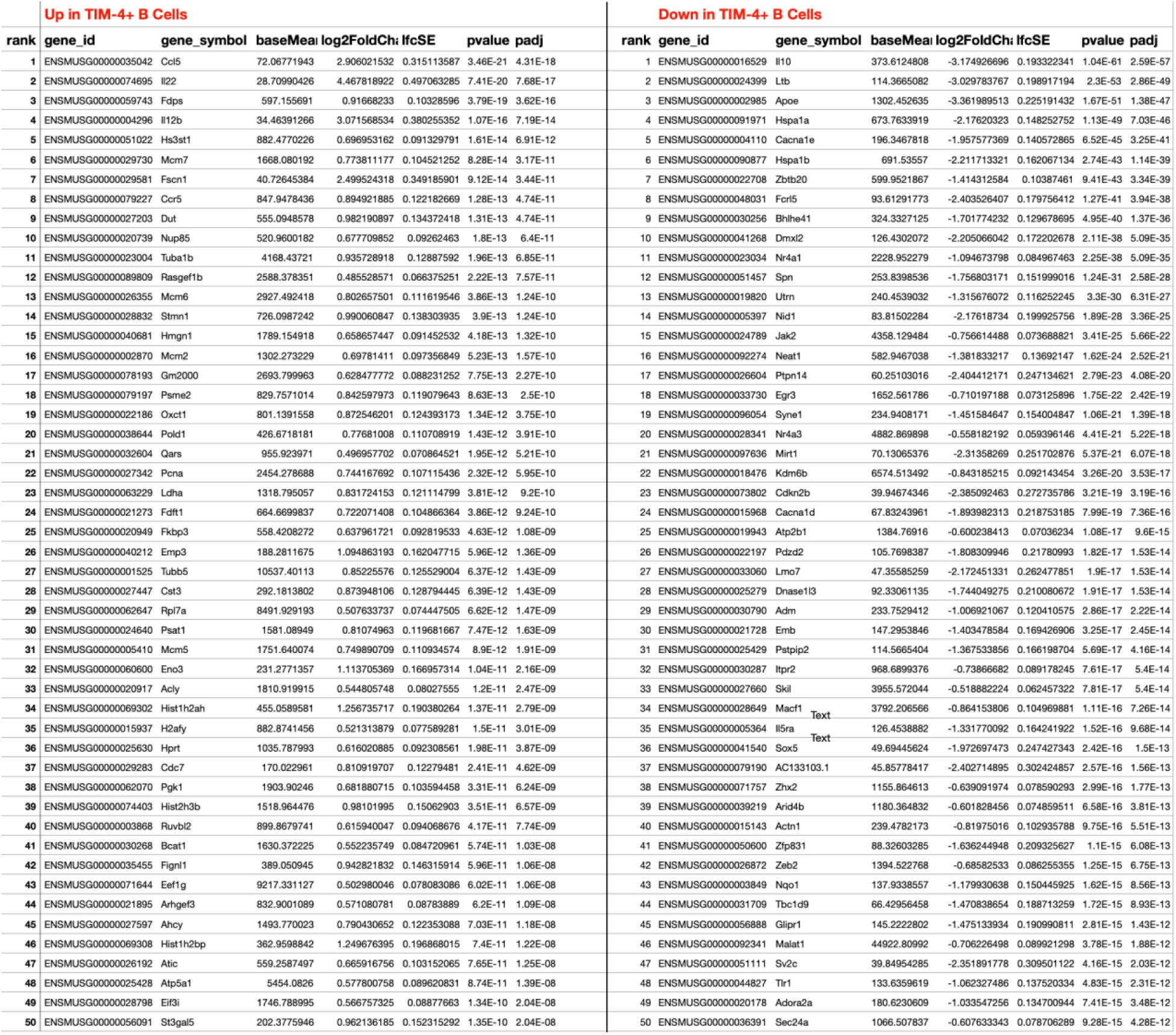
Top 50 s TIM-4+ vs. TIM-1+B cells.

**Supplementary Table 2:**
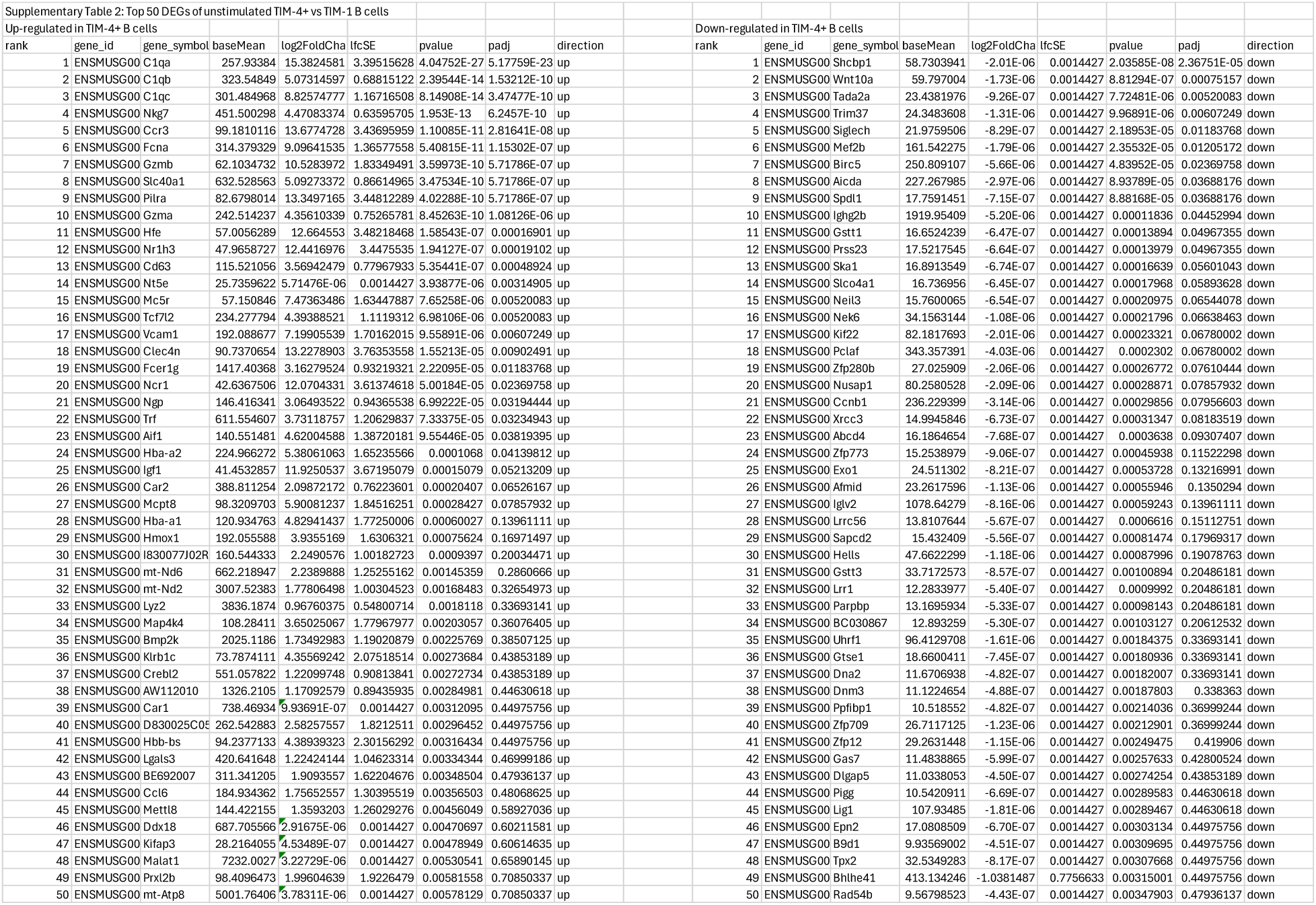
Top 50 unstimulated TIM-4+ vs. TIM-1+B cells.

## References

1. C. Mauri, M. Menon, The expanding family of regulatory B cells. International immunology 27, 479-486 (2015).

2. A. Cherukuri, Q. Ding, A. Sharma, K. Mohib, D. M. Rothstein, Regulatory and Effector B Cells: A New Path Toward Biomarkers and Therapeutic Targets to Improve Transplant Outcomes? Clin Lab Med 39, 15–29 (2019).

3. A. Cherukuri, K. Mohib, D. M. Rothstein, Regulatory B cells: TIM-1, transplant tolerance, and rejection. Immunol Rev 299, 31–44 (2021).

4. F. E. Lund, T. D. Randall, Effector and regulatory B cells: modulators of CD4+ T cell immunity. Nature reviews. Immunology 10, 236–247 (2010).

5. A. C. Lino, T. Dorner, A. Bar-Or, S. Fillatreau, Cytokine-producing B cells: a translational view on their roles in human and mouse autoimmune diseases. Immunological reviews 269, 130–144 (2016).

6. S. Fillatreau, Pathogenic functions of B cells in autoimmune diseases: IFN-gamma production joins the criminal gang. European journal of immunology 45, 966–970 (2015).

7. S. Xiao et al., Checkpoint Receptor TIGIT Expressed on Tim-1(+) B Cells Regulates Tissue Inflammation. Cell Rep 32, 107892 (2020).

8. Q. Ding et al., Regulatory B cells are identified by expression of TIM-1 and can be induced through TIM-1 ligation to promote tolerance in mice. The Journal of clinical investigation 121, 3645–3656 (2011).

9. Q. Ding, K. Mohib, V. Kuchroo, D. Rothstein, TIM-4 Identifies IFN-gamma-Expressing Proinflammatory B Effector 1 Cells That Promote Tumor and Allograft Rejection. Journal of immunology 199, 2585–2595 (2017).

10. W. Wojciechowski et al., Cytokine-producing effector B cells regulate type 2 immunity to H. polygyrus. Immunity 30, 421–433 (2009).

11. T. A. Barr et al., B cell depletion therapy ameliorates autoimmune disease through ablation of IL-6-producing B cells. The Journal of experimental medicine 209, 1001–1010 (2012).

12. S. A. Olalekan, Y. Cao, K. M. Hamel, A. Finnegan, B cells expressing IFN-gamma suppress Treg-cell differentiation and promote autoimmune experimental arthritis. European journal of immunology 45, 988–998 (2015).

13. R. A. Elsner, S. Smita, M. J. Shlomchik, IL-12 induces a B cell-intrinsic IL-12/IFNgamma feed-forward loop promoting extrafollicular B cell responses. Nat Immunol 25, 1283–1295 (2024).

14. D. A. Bermejo et al., Trypanosoma cruzi trans-sialidase initiates a program independent of the transcription factors RORgammat and Ahr that leads to IL-17 production by activated B cells. Nature immunology 14, 514–522 (2013).

15. A. Bar-Or et al., Abnormal B-cell cytokine responses a trigger of T-cell-mediated disease in MS? Ann Neurol 67, 452–461 (2010).

16. R. Li et al., Proinflammatory GM-CSF-producing B cells in multiple sclerosis and B cell depletion therapy. Sci Transl Med 7, 310ra166 (2015).

17. S. Fillatreau, C. H. Sweenie, M. J. McGeachy, D. Gray, S. M. Anderton, B cells regulate autoimmunity by provision of IL-10. Nat Immunol 3, 944–950 (2002).

18. D. Zhao et al., Innate Allorecognition and Memory in Transplantation. Front Immunol 11, 918 (2020).

19. E. J. Suchin et al., Quantifying the frequency of alloreactive T cells in vivo: new answers to an old question. Journal of immunology 166, 973–981 (2001).

20. K. I. Abou-Daya, et al., Resident memory T cells form during persistent antigen exposure leading to allograft rejection. Sci Immunol 6, (2021).

21. K. Mohib et al., Antigen-dependent interactions between regulatory B cells and T cells at the T:B border inhibit subsequent T cell interactions with DCs. Am J Transplant 20, 52–63 (2020).

22. E. Bettelli et al., Reciprocal developmental pathways for the generation of pathogenic effector TH17 and regulatory T cells. Nature 441, 235–238 (2006).

23. Y. Lee et al., Induction and molecular signature of pathogenic TH17 cells. Nat Immunol 13, 991–999 (2012).

24. C. L. Langrish et al., IL-23 drives a pathogenic T cell population that induces autoimmune inflammation. The Journal of experimental medicine 201, 233–240 (2005).

25. P. Thakore et al., The chromatin landscape of Th17 cells reveals mechanisms of diversification of regulatory and pro-inflammatory states. bioRxiv 2022.02.26.482041;, (2022).

26. S. A. Yoo et al., Placental growth factor regulates the generation of T(H)17 cells to link angiogenesis with autoimmunity. Nat Immunol 20, 1348–1359 (2019).

27. K. Shimada et al., T-Cell-Intrinsic Receptor Interacting Protein 2 Regulates Pathogenic T Helper 17 Cell Differentiation. Immunity 49, 873–885 e877 (2018).

28. B. Wu et al., RAS P21 Protein Activator 3 (RASA3) Specifically Promotes Pathogenic T Helper 17 Cell Generation by Repressing T-Helper-2-Cell-Biased Programs. Immunity 49, 886–898 e885 (2018).

29. J. T. Gaublomme et al., Single-Cell Genomics Unveils Critical Regulators of Th17 Cell Pathogenicity. Cell 163, 1400–1412 (2015).

30. Y. Bao et al., Identification of IFN-gamma-producing innate B cells. Cell Res 24, 161–176 (2014).

31. B. Stockinger, S. Omenetti, The dichotomous nature of T helper 17 cells. Nature reviews. Immunology 17, 535–544 (2017).

32. M. Pawlak, A. W. Ho, V. K. Kuchroo, Cytokines and transcription factors in the differentiation of CD4(+) T helper cell subsets and induction of tissue inflammation and autoimmunity. Current opinion in immunology 67, 57–67 (2020).

33. L. Codarri et al., RORgammat drives production of the cytokine GM-CSF in helper T cells, which is essential for the effector phase of autoimmune neuroinflammation. Nat Immunol 12, 560–567 (2011).

34. C. Wang et al., CD5L/AIM Regulates Lipid Biosynthesis and Restrains Th17 Cell Pathogenicity. Cell 163, 1413–1427 (2015).

35. G. Meyer Zu Horste et al., RBPJ Controls Development of Pathogenic Th17 Cells by Regulating IL-23 Receptor Expression. Cell Rep 16, 392–404 (2016).

36. B. U. Schraml et al., The AP-1 transcription factor Batf controls T(H)17 differentiation. Nature 460, 405–409 (2009).

37. C. C. Lin et al., Bhlhe40 controls cytokine production by T cells and is essential for pathogenicity in autoimmune neuroinflammation. Nat Commun 5, 3551 (2014).

38. L. Bod et al., Single-cell profiling identifies B-cell specific checkpoint molecules that regulate antitumor immunity. Nature 619 348–356. doi: 10.1038/s41586-023-06231-0. Epub 2023 Jun 21. PMID: 37344597, 348-356 (2023).

39. A. M. Khalil, J. C. Cambier, M. J. Shlomchik, B cell receptor signal transduction in the GC is short-circuited by high phosphatase activity. Science 336, 1178–1181 (2012).

40. E. H. Akama-Garren et al., Follicular T cells are clonally and transcriptionally distinct in B cell-driven mouse autoimmune disease. Nat Commun 12, 6687 (2021).

41. Y. Benjamini, Y. Hochberg, Controlling the False Discovery Rate: A Practical and Powerful Approach to Multiple Testing. Journal of the Royal Statistical Society. Series B (Methodological) 57, 289–300 (1995).

42. A. Zhu, J. G. Ibrahim, M. I. Love, Heavy-tailed prior distributions for sequence count data: removing the noise and preserving large differences. Bioinformatics 35, 2084–2092 (2018).

43. W. Huber et al., Orchestrating high-throughput genomic analysis with Bioconductor. Nature Methods 12, 115–121 (2015).

