## SUPPLEMENTARY METHODS for "RORγt^+^ B cells express proinflammatory cytokines and promote allo- and auto- immunity"

**Islet isolation, transplantation, and alloantigen immunization:** Allogeneic islets from B6 donors were digested with collagenase V (Sigma-Aldrich), purified by filtration through a 100-μm nylon cell strainer (BD Biosciences), hand-picked under a stereomicroscope, and placed under the left renal capsule of sex-matched allogeneic recipients with streptozotocin-induced diabetes (400 islets per recipient), as we previously described (*1, 2*). All recipients had glycemia < 150 mg/dl within 2 d after transplant. Blood glucose > 250 mg/dl for two consecutive days after engraftment was defined as rejection(*1, 2*).

**EAE**: Mice were immunized subcutaneously in the flank with an emulsion containing MOG 35–55 (200µg/mouse) and CFA (200 µl/mouse; Difco Laboratories, containing M. tuberculosis H37Ra, 5 mg/ml). Pertussis toxin (200 ng/mouse; List Biological Laboratories) was administered intraperitoneally on days 0 and 2. Mice were monitored and assigned grades for clinical signs of EAE. Mice were observed for signs of EAE beginning on day 7 after immunization. Mice were clinically assessed with daily assignment of scores on a standard 0–5 scale as follows: no clinical expression of disease, 0; partially limp tail, 0.5; completely limp tail, 1; limp tail and waddling gait, 1.5; paralysis of one hind limb, 2; paralysis of one hind limb and partial paralysis of the other hind limb, 2.5; complete paralysis of both hind limbs, 3; ascending paralysis, 3.5; paralysis of trunk, 4; moribund, 4.5; death, 5. To evaluate the suppressive ability of IL-17A^-/-^ B cells, purified total IL-17A^-/-^ B cells, B cells were transferred i.v. into µMT recipients. Hosts were then immunized to induce EAE.

**Adoptive transfer of TCR-transgenic T cells:** CD4 and CD8 T cells from spleens and lymph nodes of OT-II and OT-I transgenic mice were purified using EasySep™ Mouse CD4^+^ and CD8^+^ T cell Isolation Kits (STEMCELL), respectively, and were labeled with CFSE (ThermoFisher). and 2×10^6^ cells were injected i.v. per recipient. Mice were immunized the next day with splenocytes from Act-mOVA F1 mice. Mice were sacrificed and transferred OT-I or OT-II cells analyzed at 4 days for proliferation (CFSE dilution), and at 7 days for cytokine expression (flow cytometry).

**Gene set ordinal association test:** To identify biological processes associated with differentially expressed genes, gene set enrichment was performed using the gene set ordinal association test (GOAT). Gene sets were obtained from the mSigDB Hallmark and Gene Ontology gene sets (via msigdbr v25.1.1) (*3, 4*). GOAT was performed using the naïve bootstrapping approach as implemented in the *goat* R package and as described previously (*5*). Briefly, GOAT was performed separately for pairwise comparison between stimulated or unstimulated TIM-4^+^, TIM-1^+^, or DN B cells, using the differential gene expression results from DESeq2. The shrunken fold changes and adjusted p values from DESeq2 were used as input for all genes in each comparison, and gene sets with FDR-adjusted p-values ≤ 0.05 after the two-step multiple testing correction implemented in the *goat* R package (v 1.1.2) were retained for downstream analysis and interpretation.

**References:**

1.	Q. Ding, K. Mohib, V. Kuchroo, D. Rothstein, TIM-4 Identifies IFN-gamma-Expressing Proinflammatory B Effector 1 Cells That Promote Tumor and Allograft Rejection. *Journal of immunology* **199**, 2585-2595 (2017).

2.	Q. Ding *et al.*, Regulatory B cells are identified by expression of TIM-1 and can be induced through TIM-1 ligation to promote tolerance in mice. *The Journal of clinical investigation* **121**, 3645-3656 (2011).

3.	A. Liberzon *et al.*, The Molecular Signatures Database (MSigDB) hallmark gene set collection. *Cell Syst* **1**, 417-425 (2015).

4.	M. Ashburner *et al.*, Gene ontology: tool for the unification of biology. The Gene Ontology Consortium. *Nat Genet* **25**, 25-29 (2000).

5.	F. Koopmans, GOAT: efficient and robust identification of gene set enrichment. *Commun Biol* **7**, 744 (2024).
